# *BioPKS Pipeline*: An integrated platform for merging the computational design of chimeric type I polyketide synthases with enzymatic pathways for chemical biosynthesis

**DOI:** 10.1101/2024.11.04.621673

**Authors:** Yash Chainani, Jacob Diaz, Margaret Guilarte-Silva, Vincent Blay, Quan Zhang, William Sprague, Keith E. J. Tyo, Linda J. Broadbelt, Aindrila Mukhopadhyay, Jay D. Keasling, Hector Garcia Martin, Tyler W. H. Backman

**Affiliations:** Department of Chemical and Biological Engineering, Northwestern University, Evanston, IL, USA; Center for Synthetic Biology, Evanston, IL, USA; Joint BioEnergy Institute, Emeryville, CA, USA; Biological Systems and Engineering Division, Lawrence Berkeley National Laboratory, Berkeley, CA, USA; QB3, University of California, Berkeley, Berkeley, CA, USA; Department of Chemical and Biomolecular Engineering, University of California, Berkeley, Berkeley, CA, USA; Department of Bioengineering, University of California, Berkeley, Berkeley, CA, USA; Center for Biosustainability, Danish Technical University, Lynby, Denmark; BCAM, Basque Center for Applied Mathematics, Bilbao, Spain; DOE Agile BioFoundry, Emeryville, California

## Abstract

Synthetic biology offers the promise of manufacturing chemicals more sustainably than petrochemistry. Yet, both the rate at which biomanufacturing can synthesize these molecules and the net chemical accessible space are limited by existing pathway discovery methods which can often rely on arduous literature searches. Here, we present an automated retrobiosynthesis tool, BioPKS Pipeline, that simultaneously tackles both problems by integrating multifunctional type I polyketide synthases (PKSs) with monofunctional enzymes to propose the synthesis of desired target chemicals via two new tools: DORAnet and RetroTide. While monofunctional enzymes are valuable for carefully decorating a substrate’s carbon backbone, they typically cannot expand the backbone itself. PKSs can, instead, predictably do this through their unique ability to catalyze carbon-carbon bond formation reactions iteratively. We have evaluated the performance of BioPKS Pipeline against a previously published set of 155 molecules of interest for biomanufacturing, and report that BioPKS Pipeline could produce exact designs for 93 of them, as well as pipelines to a chemically similar product for most of the remaining molecules. Furthermore, BioPKS Pipeline successfully proposes biosynthetic routes for complex therapeutic natural products (cryptofolione and basidalin) for which no known biosynthetic pathway currently exists.

## Introduction

An increasing concern regarding climate change has led to growing public and private interest in synthetic biology to enable the biomanufacturing of a wide array of commodity chemicals^1–5^. In 2022, the United States’ government signed an executive order to invest 2 billion dollars into the U.S. bioeconomy to divert chemical manufacturing away from pollutive fossil-fuel based processes and towards environmentally benign bioengineered ones^6^. Further, the overall demand for renewable products has also been solidifying, driven by the need to achieve sustainability in manufacturing while reducing carbon emissions to mitigate rising global temperatures^7,8^.

Despite this increasing interest, the design of biosynthetic pathways to produce key commodity chemicals without known biosynthesis pathways remains a bottleneck^9–11^. Long and cumbersome literature searches can be common in piecing together known or potentially promiscuous enzymatic reactions that may eventually lead to a target product from simple precursors and intermediates. Such manual pathway discovery methods diminish the rate at which biomanufacturing can replace the ∼3500 or so high-volume, petroleum-derived chemicals currently in circulation^12,13^.

In addition to this slow rate of access, the net chemical space that existing methods can reach is also limited by the customarily exclusive use of monofunctional enzymes. Monofunctional enzymes catalyze individual reactions only and, to date, comprise a majority of engineered pathways^4,14,15^. While such enzymes are valuable for the precise modification of functional groups on a substrate’s carbon backbone, they offer few chances to increase the length of the backbone itself. Consequently, relying exclusively on monofunctional enzymes often restricts the achievable carbon chain length of target products to about that of the original carbon source. Metabolic maps of bio-based chemicals in the literature highlight that pathways beginning from pyruvate and other central carbon metabolites yield C3-C8 products only^16,17^. Such short backbones render it difficult for synthetic biology to compete with the longer hydrocarbons (>C10) present in petrochemical feedstocks^18^, especially when making products that are more structurally complex than linear carbon chains.

In fact, attempting to synthesize molecules solely using monofunctional enzymes overlooks how nature synthesizes many valuable and complex natural products: by combining multifunctional and monofunctional enzymes for complementary purposes. Multifunctional enzymes (which catalyze multiple reactions) are used by these natural pathways, genetically encoded in the biosynthetic gene clusters (BGCs) of several microbes, plants, and fungi, to first construct elongated carbon scaffolds. Only after the construction of such scaffolds are monofunctional enzymes used for structural fine-tuning^19–22^. A particularly interesting and valuable class of multifunctional enzymes are polyketide synthases (PKSs), which can catalyze the expansion of carbon backbones^23–25^. PKSs are responsible for the synthesis of many medicinally relevant and structurally diverse molecules, from macrocyclic antibiotics (e.g., erythromycin^26^) to long aliphatic anticancer therapeutics (e.g., gephyronic acid^27,28^). PKSs broadly fall into one of three categories: type I, II, and III^23^. Type II and III PKSs consist of discrete, unlinked enzymes operating independently^29,30^ while type I PKSs comprise several covalently linked enzymatic domains working together as an assembly-line to synthesize large polyketides from simple building blocks^31^. These building blocks are typically acyl-coenzyme A (acyl-CoA) derivatives, such as acetyl-CoA or malonyl-CoA^23,25^. Through decarboxylative Claisen condensation reactions performed by their ketosynthase (KS) domain, PKSs are able to iteratively add 2 to 3 carbon atoms from such acyl-CoA units onto a previously loaded acyl-CoA substrate^23,25^. Over the course of several modules, each of which minimally contains a KS, an acyltransferase (AT), and an acyl carrier protein (ACP) domain, PKSs can extend the initial substrate from just 2 to 6 carbon atoms to over 20 in length^23,25^. This unique ability of PKSs to catalyze C-C bond formation reactions is immensely beneficial for biomanufacturing and could be repurposed towards the synthesis of chemicals, such as biofuels, for which carbon chain length correlates positively with fuel efficiency^32^.

Beyond constructing elongated backbones, PKSs also allow for the incorporation of diverse side chains and varying the level of reduction of the synthesized molecule^33–38^. While AT domains primarily function to extend the carbon chain arising from the previous module by selecting common acyl-CoA units, such as malonyl-CoA or methylmalonyl-CoA, there is a large diversity of *α*-substituted starter and extender acyl-CoA units that can be selected by AT domains^38–40^. For instance, when designing monomers, terminal alkene groups (useful for polymerization) may be included through unsaturated units such as allylmalonyl-CoA^41,42^. Meanwhile, aromatic rings, a key structural component of many monomers and polymers, can be added via aromatic units such as cinnamoyl-CoA^43^. Recent advances in engineering strategies for AT domain swapping^44,45^ have enabled the incorporation of such atypical acyl-CoA units into polyketide products. In particular, the use of the highly promiscuous mammalian malonyl acetyltransferase (MAT) has further broadened the range of possible extender units for PKS engineering^46^. Alongside performing AT domain swaps, the oxidation state and functionalization of each extender unit added to the growing carbon backbone can be tuned further by swapping or modifying the PKS reductive loop^47–49^, which comprises the ketoreductase (KR), dehydratase (DH), and enoylreductase (ER) domains. Moreover, since a PKS substrate is always tethered to its synthase, the likelihood of intermediate loss can be lower with PKSs than in engineered metabolic pathways wherein intermediates may be funneled into different pathways altogether while being shuttled around individual enzymes in the cell^50^. Such derailments can lead to unexpected^51–56^, or even toxic^57^ byproducts. Given the predictable deterministic logic of Type I PKSs, however, one can expect the final product to be directly correlated with the domains and acyl-CoA units involved^26^.

This modular and predictable, yet tunable, nature of multifunctional PKSs makes them an attractive platform for biomanufacturing. While PKSs have historically been challenging to engineer and are characterized by low turnover rates^33,45^, ongoing research has only further confirmed their capacity for structural derivatization: recently, several researchers have engineered the deoxyerythronolide synthase (DEBS) involved in erythromycin’s biosynthesis to accept fluorinated acyl-CoA units so as to regioselectively incorporate fluorine into the erythromycin scaffold^58,59^. This has also been achieved with other PKS products as well^60^, which is crucial since an increasing number of approved drugs today comprise fluorine atoms^61,62^. Because PKSs follow a modular deterministic logic, an engineered PKS provides a straightforward path to create a large library of chemical analogues that can be explored for improved properties via domain and module engineering^44^. We believe that combining the versatility and scaffolding ability of PKSs with the precision of monofunctional enzymes, thereby mimicking the biosynthesis of natural products, can ultimately unlock a wider space of chemicals for biomanufacturing than would be accessible with either type of enzyme alone.

Here, we present BioPKS Pipeline, a novel and automated retrobiosynthesis software that seamlessly integrates the design of multifunctional chimeric type I PKSs with that of monofunctional enzymatic pathways into a single platform (Fig. 1). Given the vast design space that is already inherent to both chimeric PKSs and metabolic pathways comprising monofunctional enzymes, the combination of these unlocks a design space that is considerably larger than either route alone. To efficiently navigate this larger space, BioPKS Pipeline first uses RetroTide, a chimeric PKS tool presented here to construct the carbon scaffold of a target molecule, and subsequently, DORAnet to tailor the PKS product towards the final target through monofunctional enzymes. While several *in silico* retrobiosynthesis tools already exist for monofunctional enzymes^10,11,63–75^, to our knowledge, none exist for designing chimeric multifunctional enzymes, and especially not for the combination of the two. Existing tools^76,77^ and databases^78–80^ for multifunctional enzymes are valuable in matching known natural product structures to known biosynthetic gene clusters, but do not suggest novel PKS chimeras nor post-PKS modification pathways for a given target structure (that may or may not classify as a natural product). If an integrated pathway to a desired target exists, BioPKS Pipeline provides: (1) a chimeric PKS design to synthesize a PKS intermediate product as structurally similar to the target as possible, and (2) a monofunctional enzymatic pathway design to transform this PKS intermediate into the final target. Additional capabilities of utility to the synthetic biologist have also been included within BioPKS Pipeline for in-depth post-PKS pathway analyses. These include reaction feasibility prediction and thermodynamics calculations^81,82^.

**Fig. 1:**
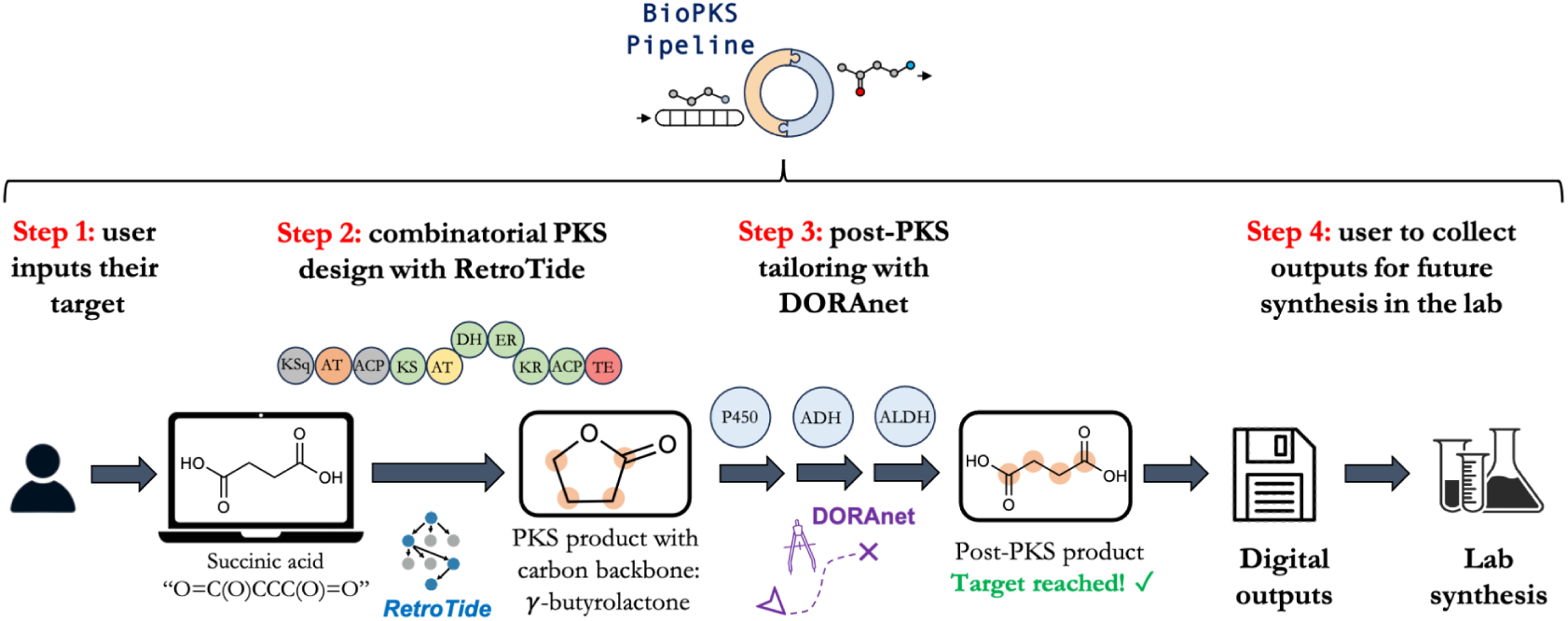
BioPKS Pipeline reaches target molecules *in silico* by first constructing their carbon backbone and then performing structural refinements. BioPKS Pipeline uses both multifunctional type I polyketide synthases (PKSs) and monofunctional enzymes for chemical biosynthesis. Our platform is built upon two key rule-based components – RetroTide and DORAnet. When a user inputs a target structure, RetroTide first suggests chimeric PKS designs to synthesize the carbon backbone of this target. DORAnet then performs enzymatic post-PKS decorations of the PKS product to reach the target chemical. If an integrated PKS and post-PKS pathway design is found, the user can collect this design from BioPKS Pipeline for future experimental synthesis in the laboratory.

We have demonstrated the capabilities of BioPKS Pipeline to recommend biosynthetic pathways across three distinct scenarios: a structurally diverse set of 15 molecules, 155 targets of relevant biomanufacturing candidates, and two natural therapeutic products for which no biosynthetic pathway is currently known. First, we deployed our tool to suggest biosynthetic pathways to 15 structurally diverse molecules of varying carbon chain lengths (from 3 to 18) that we curated. In doing so, only a single post-PKS modification step was allowed and, for all 15 molecules, BioPKS Pipeline was able to construct the requisite carbon backbone. Subsequently we validated BioPKS Pipeline against a larger set of 155 molecules obtained from a previously published list of potential biomanufacturing candidates^13^, allowing for up to 2 post-PKS modification steps this time. Against this set, BioPKS Pipeline was able to suggest pathways to completely synthesize 60% of molecules. As a final test on the general capabilities of our tool, we prompted BioPKS Pipeline to synthesize the therapeutic natural products cryptofolione^83^ and basidalin^84^, for which no known biosynthetic pathways currently exist. Remarkably, BioPKS Pipeline succeeded in suggesting PKS designs and post-PKS pathways to synthesize these products completely.

Altogether, our work represents one of the first *in silico* demonstrations of combining the power of PKSs to expand carbon backbones with the capability of monofunctional enzymes to engineer a broad range of molecules - from simple commodities to complex natural products.

## Results

### Architecture of BioPKS Pipeline

BioPKS Pipeline is an automated retrobiosynthesis tool with two key rule-based components (Fig. 2): RetroTide (presented here) and DORAnet (publicly available at https://github.com/wsprague-nu/doranet). When a target chemical is input, BioPKS Pipeline first attempts to synthesize this molecule and its carbon skeleton using only PKSs via RetroTide, which in turn explores different combinations of known PKS domains and organizes them into modules. Each domain has a single reaction rule associated with it based on its known enzymatic function. Over the course of a single module, the application of reaction rules associated with each constituent domain always results in the carbon-chain elongation of the product arising from the previous module as well as any further decorations due to the presence of either the KR, DH, and/ or ER domains. PKS designs ranked by the chemical similarity of the corresponding PKS product in comparison to the target are then provided by RetroTide (see Methods for the different similarity metrics available to users). If a target can be synthesized by PKSs alone, BioPKS Pipeline terminates and users can search our previously released ClusterCAD 2.0 database^85^ (https://clustercad.jbei.org/pks/) for natural PKS designs closely matched with the predicted chimera. This search will indicate the genetic edits, such as domain swaps, insertions, and/or truncations needed to produce the chimera from the closest matched natural PKS and the BGCs encoding for such a PKS. ClusterCAD 2.0^85^ is a database of naturally-occurring type I PKSs, non-ribosomal peptide synthetases (NRPSs), as well as PKS-NRPS hybrids, annotated and searchable to guide selection of parts for PKS engineering. Crucially, ClusterCAD 2.0 supports domain architecture, chemical structure, as well as protein sequence searches such that users can extract the most similar natural megasynthase to their query in terms of either domain architecture, chemical similarity, or sequence similarity.

**Fig. 2:**
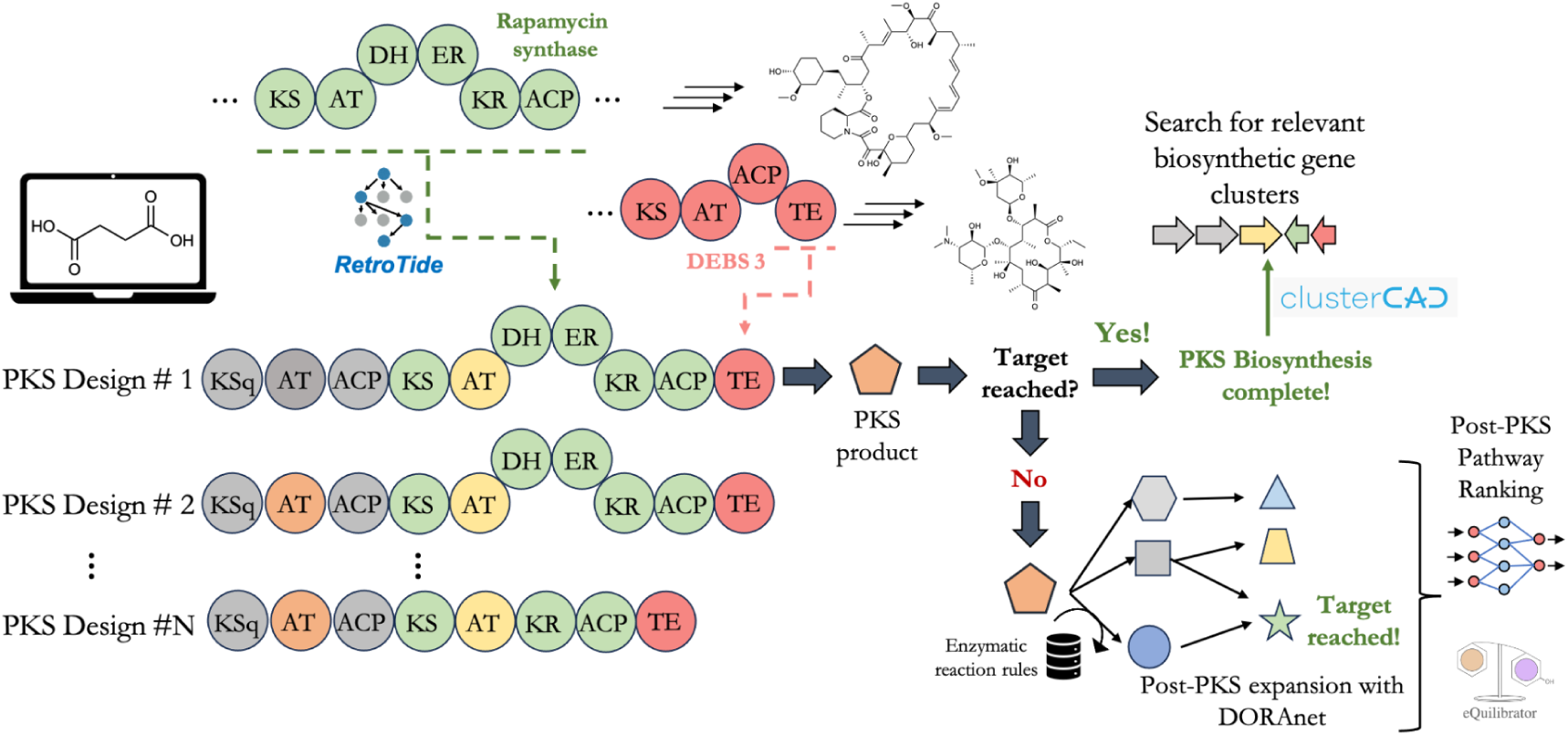
BioPKS Pipeline combinatorially designs novel, chimeric PKSs from existing PKSs to produce intermediates structurally similar to the target molecule. When a target molecule is input, BioPKS Pipeline first calls upon RetroTide to suggest chimeric PKS designs by combinatorially mixing domains from existing PKSs, such as rapamycin synthase or 6-deoxyerythronolide B synthase (DEBS). Here, each color represents a domain from a distinct known PKS. RetroTide’s PKS designs are output in descending order of chemical similarity as measured between the corresponding PKS product and the target chemical. If the target can be synthesized solely with PKSs, BioPKS Pipeline stops and users can search the ClusterCAD 2.0 database for closely matched PKSs. If the target cannot be synthesized by PKSs, then BioPKS Pipeline calls upon DORAnet to modify the top-ranked PKS product with monofunctional enzymes, whose activities are digitally encoded as reaction rules. The post-PKS pathways generated can then be ranked with thermodynamics and/ or pathway feasibility analyses. If the top-ranked PKS product does not lead to the final target through post-PKS modifications, then alternate PKS designs are considered.

If the desired target cannot be produced by PKSs alone, however, then enzymatic post-PKS modifications via DORAnet are attempted on the PKS product that is most chemically similar to the target molecule.

DORAnet is a metabolic network expansion tool that then performs these post-PKS modifications by recursively expanding upon the PKS product from RetroTide using our previously published reaction rules, which encode for the potential promiscuities of known enzymes^86,87^. Each reaction rule within DORAnet is associated with a list of enzymes that follow the same reaction pattern. When creating reaction networks with DORAnet, users can choose between either our set of 1224 generalized reaction rules (JN1224MIN^86^), or our set of 3604 intermediate rules (JN3604IMT^87^). Our generalized rules assume a high level of promiscuity across all enzymes by focusing only on the potential reaction centers within substrates that could directly participate in a predicted reaction^86^. Meanwhile, our intermediate rules assume a lower level of promiscuity by considering not only such reaction centers within substrates, but also the surrounding chemical context within which these reaction centers exist^87^. Both of our rulesets were extracted from the publicly available MetaCyc database^88^ and more information on their coverage of biochemistry reported within other databases^89,90^ can be found in the Methods section (“Performing post-PKS modifications on RetroTide products with DORAnet”). The recursive application of reaction rules by both RetroTide and DORAnet as well as the passing of the polyketide product output by RetroTide into DORAnet is completely automated by BioPKS Pipeline and does not require any manual intervention from the user.

Regardless of the reaction ruleset chosen by the user, depending on the number of post-PKS modification steps specified, DORAnet may enumerate far more post-PKS pathways than can practically be manually analyzed. Thus, to elucidate the most feasible pathway chemistries for experimentation, our previously released DORA-XGB^91^ reaction feasibility classifier has been incorporated into BioPKS Pipeline to rank post-PKS reactions by their feasibility. DORA-XGB^91^ is publicly available at https://github.com/tyo-nu/DORA_XGB, and scores the feasibility of a given enzymatic reaction between 0 and 1, wherein a higher score represents a more feasible reaction. DORA-XGB was trained by first dividing reported reactions from publicly available metabolic databases into sets of thermodynamically downhill (positive) and uphill (negative) reactions. From the set of downhill reactions, even more negative reactions were synthetically generated by considering alternate reaction sites that an enzyme could have transformed but were still not experimentally observed to undergo catalysis. Consequently, DORA-XGB’s score reflects both the thermodynamic feasibility of a reaction under typical physiological conditions in the cell as well as the likelihood that a specific functional group on a substrate would undergo catalysis. A lower scoring DORAnet reaction then indicates either that a proposed reaction is thermodynamically uphill or that the suggested reaction site is unlikely to undergo catalysis by known enzymes, signifying the need for some form of enzyme engineering.

In the event that post-PKS modifications of the top PKS product cannot yield the final target molecule, BioPKS Pipeline pursues two alternatives to further attempt to synthesize this product. Firstly, alternate, lower-ranked PKS designs and their corresponding PKS products from RetroTide are considered (Fig. 2). This is because a structurally dissimilar PKS intermediate may actually prove to be more amenable to the enzymatic modifications required to reach the final target molecule. The maximum number of alternate PKS designs considered is a hyperparameter that can be tuned by users, but is set to 5 by default to limit computational expense. The second alternative involves working backwards from the desired chemical target with DORAnet to identify alternative precursors that can lead to this target. These intermediates can then be tested with RetroTide to see if they can be synthesized exactly by a PKS. This alternative approach can identify PKS precursors initially rejected by RetroTide, due to low overall chemical similarity to the final product. These contingencies and our primary algorithm driving BioPKS Pipeline ensure that the space of combined pathways has been adequately explored. The default configurations for BioPKS Pipeline produce results in just minutes on standard personal computers, but can be scaled to explore a larger chemical space on more powerful computer systems.

### Selection of starter and extender acyl-CoA derivatives by RetroTide

Acyl-CoA derivatives are the key building blocks through which PKSs synthesize polyketide products (Fig. 3). All type I PKSs begin their assembly line synthesis of polyketides by first loading an acyl-CoA or acyl-ACP derivative via their loading module. In this work, we refer to this initial acyl-CoA unit as a starter unit while an acyl-CoA used by a downstream extension module is referred to as an extender unit. The loading module of a given PKS can often contain a ketosynthase-like (KSq) domain, which if present, loads a dicarboxylic acid wherein one of the acid groups has been acylated by CoA (e.g., malonyl-CoA). If this KSq domain is absent in the loading module, then PKSs load a monocarboxylic acid wherein the single acid group has already been CoA-acylated (e.g., propionyl-CoA or acetyl-CoA).

**Fig. 3:**
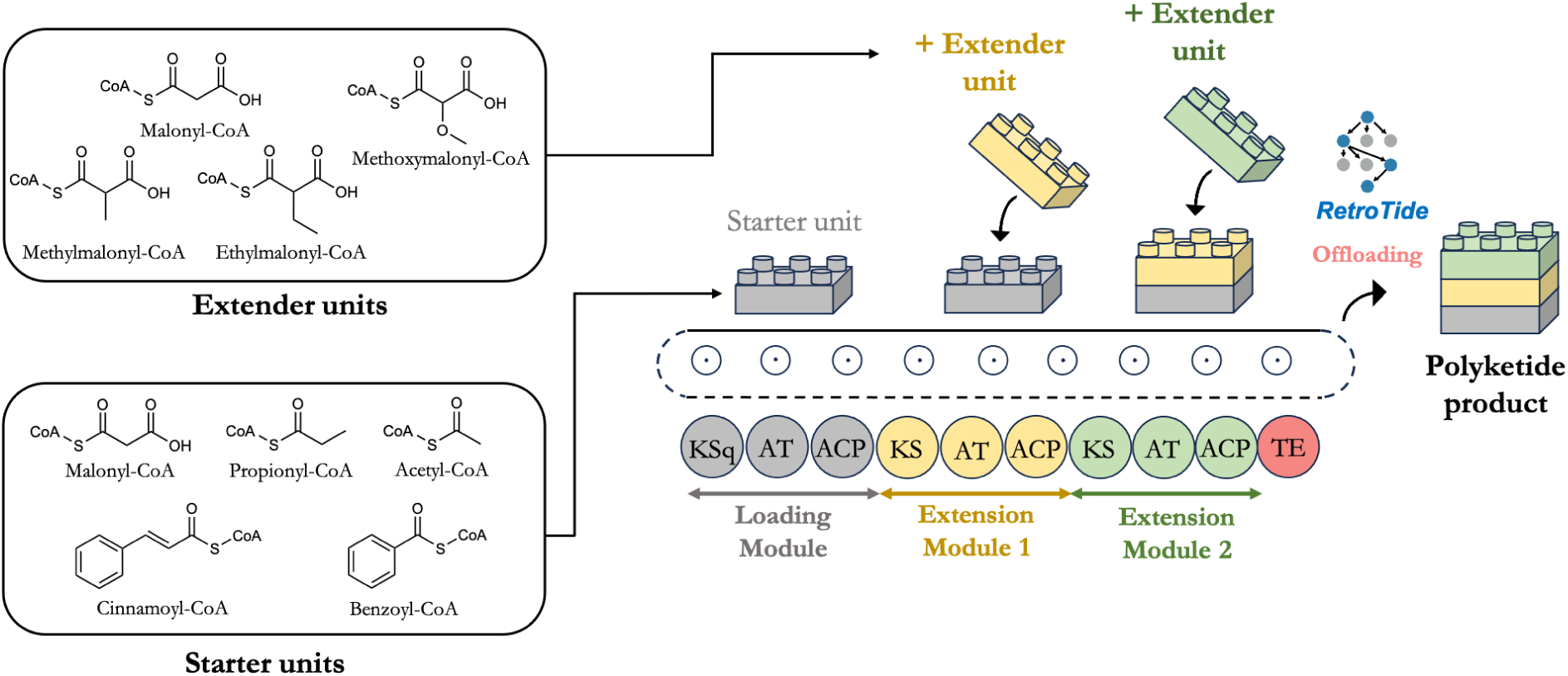
RetroTide builds polyketide products *in silico* using a large and customizable set of starter and extender acyl-CoA derivatives. PKSs use acyl-CoA derivatives as building blocks for synthesizing complex and functionally diverse polyketides. The loading module of PKSs can often contain a ketosynthase-like (KSq) domain, which if present, loads dicarboxylic acids wherein one of the acid groups has been acylated by CoA (e.g., malonyl-CoA). If this KSq domain is absent, however, then PKSs load monocarboxylic acids wherein the single acid group has already been acylated by CoA (e.g., propionyl-CoA or acetyl-CoA). Meanwhile, the ketosynthase (KS) domains in downstream extension modules use only CoA-acylated dicarboxylic acids as extension units. By considering a large and customizable set of acyl-CoA derivatives as starter and extender units within these constraints, RetroTide can build polyketide products that are as structurally similar to the final target product as possible.

In order to compile a list of starter units for RetroTide, starter units from canonical type I PKSs documented within the ClusterCAD 2.0^85^ database were first considered. This list was then expanded further with more unusual starter units that have either previously been shown to be type I PKS extenders that could be converted into starters via domain or module engineering^92^. Certain starter units from type II or type III PKSs were also included so as to expand the chemical space of molecules that can be reached. The complete list of 30 starter units available within RetroTide and their SMILES strings are reported in SI Table 1. We aimed to include an expansive set of rare and hypothetical starters (30 total) and extenders (11 total; SI Table 2) that can be pared down by users as appropriate, rather than limiting the chemical space by only including the most common PKS substrates. Note that in these starter and extender chemical structures, [S] is used as a placeholder for the substrates attachment to a CoA or ACP. Many substrates may exist in both ACP and CoA forms, and are included only once in RetroTide, but the resulting designs can be interpreted as suitable for either substrate. RetroTide also does not simulate KSq decarboxylation, so to include designs which use both carboxylated and decarboxylated starters, both variants must be included in the starter database.

We emphasize that users also have the option to either add to or trim the list of starters and extenders that Retrotide has access to depending on the experimental feasibility of working with a given starter or extender in the laboratory. Constraining the list of starters and extenders can enable BioPKS Pipeline to converge to a pathway design faster for a given target but may be accompanied by the tradeoff of reaching a smaller chemical space. Further, RetroTide builds chimeric PKSs using well-characterized domains with known enzymatic function - namely, the KR, DH, ER, AT, ACP, and TE domains - but the specific combinations of domains within a module, as well as the starter or extender substrate used, may be novel. This is intentional because it enables RetroTide to not only recapitulate existing PKS modules found in nature but to also suggest novel ones that may be possible to engineer. We recognize that not all of these suggested designs may necessarily be experimentally viable but given recent advances in PKS engineering^44,45,47–49,60,93^, we thought it best to explore as wide a space of polyketides as possible.

### Showcasing BioPKS Pipeline through the biosynthesis of two simple molecules

We report here two use-cases that exemplify, with simple molecules, the two complementary approaches used in BioPKS Pipeline (Fig. 4). The first involves the biosynthesis of 4-hydroxybutyric acid, a key intermediate in the downstream production of various C4 chemicals and for which bioengineered pathways currently exist^94–96^.

**Fig. 4:**
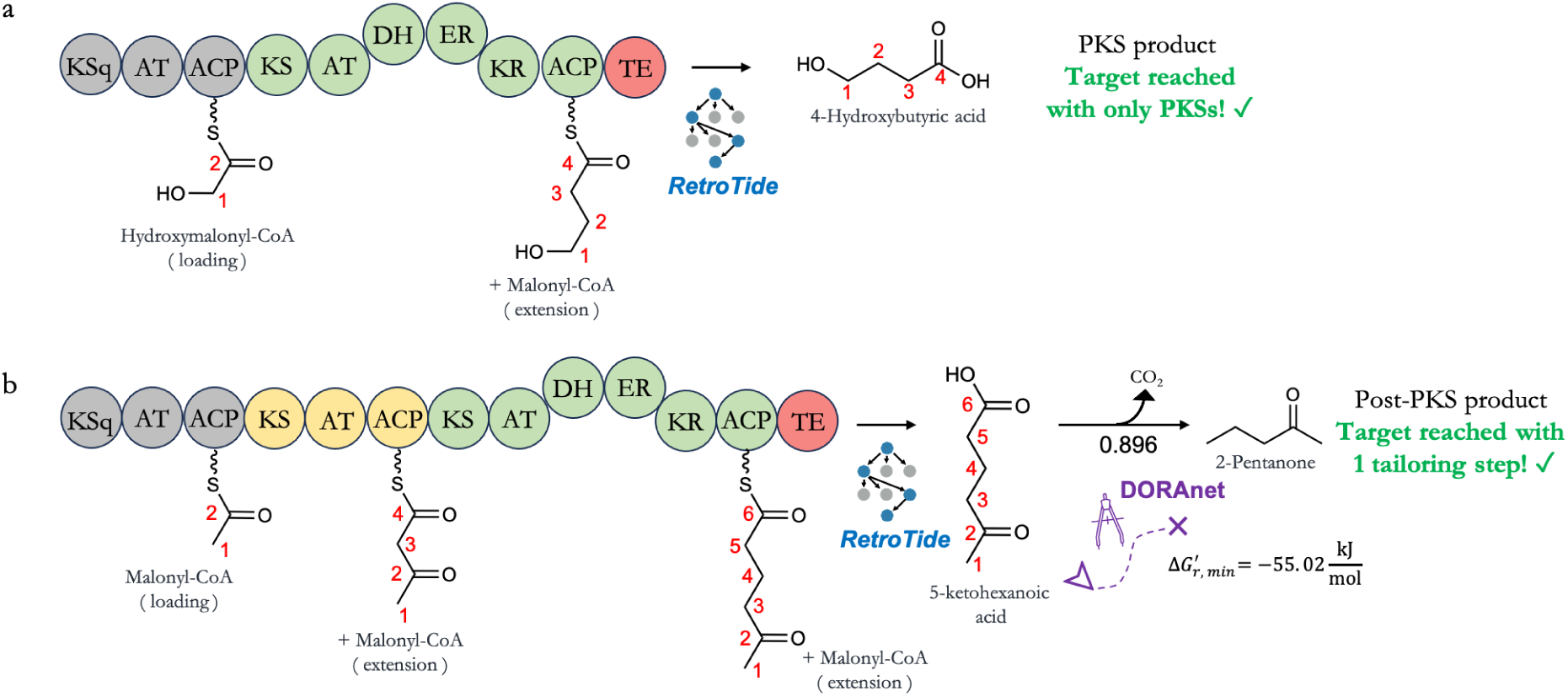
Demonstrated biosynthesis of 4-hydroxybutyric acid and 2-pentanone with BioPKS Pipeline. **a** When 4-hydroxybutyric acid is input as a target structure into BioPKS Pipeline, the top PKS design from RetroTide synthesizes 4-hydroxybutyric acid completely in just two modules. Hydroxymalonyl-CoA is recommended as a starter unit while malonyl-CoA is recommended as an extender unit. Since any additional post-PKS modifications are not required, BioPKS Pipeline terminates. **b** Conversely, when 2-pentanone is input as a target structure, the top PKS design from RetroTide reaches only as far as 5-ketohexanoic acid. DORAnet subsequently takes over and suggests a thermodynamically favorable decarboxylation reaction to synthesize 2-pentanone. This decarboxylation step is also predicted to be highly feasible with an enzymatic reaction feasibility score of 0.896.

When 4-hydroxybutyric acid is input as a target into BioPKS Pipeline, the top PKS design from RetroTide synthesizes 4-hydroxybutyric acid completely (Fig. 4A). This predicted chimera features one loading module and one extension module. RetroTide suggests that the loading module should load hydroxymalonyl-CoA as a starter unit to account for the terminal hydroxyl group in 4-hydroxybutyric acid while the extension module should load malonyl-CoA as an extender unit to complete the C4 backbone of 4-hydroxybutyric acid. Note that this may not be a naturally occurring PKS starter, please see the section “Selection of starter and extender acyl-CoA derivatives by RetroTide” for our rationale in including it anyways. The domain architecture of this extension module features a fully reducing DH-ER-KR loop that reduces the acetyl group in hydroxymalonyl-CoA to produce the fully reduced C4 backbone of 4-hydroxybutyric acid. Since the target molecule is reached here with the top PKS design, BioPKS Pipeline terminates and does not need to attempt any post-PKS modifications.

When 2-pentanone is input as a target into BioPKS Pipeline, however, the top-ranked PKS design only takes the user as far as 5-ketohexanoic acid, and needs completion through the use of DORAnet (Fig. 4B). The PKS design involves two extension modules in addition to the loading module, and malonyl-CoA is used throughout as both a starter and an extender unit. Since the final 2-pentanone target has not been reached, BioPKS Pipeline attempts to modify the 5-ketohexanoic acid PKS product by calling on DORAnet. A thermodynamically downhill decarboxylation reaction is subsequently suggested for transforming 5-ketohexanoic acid into 2-pentanone. We have used the open-source package eQuilibrator 3.0^81^ to compute reaction thermodynamics here, but the code for BioPKS Pipeline has been modularized such that other open-source tools, such as dGPredictor^82^, may easily be used as well. The suggested decarboxylation reaction also receives a high predicted feasibility score of 0.896 by DORA-XGB^91^. Searching our intermediate reaction rules database for potentially promiscuous enzymes that may be able to catalyze this reaction reveals 39 enzymes associated with the rule governing this decarboxylation. By comparing the chemical similarity of each suggested enzyme’s native substrate to that of 5-ketohexanoic acid, the enzyme acetoacetate decarboxylase emerges as a promising candidate (SI Fig. 1). This enzyme has also previously been shown to be sufficiently promiscuous to decarboxylate levulinic acid to produce 2-butanone^97^. Levulinic acid shares a similar structure to 5-ketohexanoic acid with just one less carbon atom present in its backbone between the ketone and acid groups. Alternatively, another PKS design suggested by RetroTide produces 3-ketohexanoic acid. In this PKS design, the fully reducing DH-ER-KR loop appears in the first extension module while the second extension module features only the KS, AT, and ACP domains. Given that methyl ketone synthases have previously demonstrated decarboxylation activity on keto-acids^98,99^, 3-ketohexanoic acid may also be a useful intermediate in the synthesis of 2-pentanone.

In these two simple use cases, PKSs are shown to be valuable in synthesizing the carbon backbones of user-specified targets from simple acyl-CoA derivatives while post-PKS modifications are useful for structural refinements of the PKS product to reach the final target. For more complex targets, users may wish to allow longer post-PKS pathways, though this may sometimes be accompanied with the tradeoff of potentially more infeasible or thermodynamically bottlenecked post-PKS reactions. If post-PKS pathways spanning more than one reaction step are used, then BioPKS Pipeline ranks post-PKS pathways generated by DORAnet on the basis of each pathway’s net feasibility score. This net score is in turn computed by taking the product of each constituent reaction’s DORA-XGB feasibility score within a single pathway.

### Testing BioPKS Pipeline against a curated set of fifteen structurally diverse molecules

In order to ascertain the general utility of PKSs in synthesizing a variety of carbon backbones and to investigate the utility of considering lower-ranked PKS designs, we prototyped BioPKS Pipeline against 15 molecules that we curated for their structural diversity (Fig. 5). These consist of 2 cyclic (caprolactone and γ-butyrolactone) and 13 aliphatic (tiglic acid, acrylic acid, levulinic acid, adipic acid, heptane, aminobutyric acid, 1,4-butanediol, butanone, maleic acid, muconic acid, nonadecene, sphingosine, and limonene) molecules with various functional groups. While several of these compounds have known biosynthetic pathways, we selected them as a test to evaluate our system’s ability to both create new PKS based pathways, and reproduce known PKS based pathways. For each of these 15 molecules, 5 alternate PKS designs were considered beyond the top-ranked PKS design. For testing, we constrained the number of alternate PKS designs to 5 to limit the computational complexity of networks that may be generated with each post-PKS product. This, however, is a user selectable parameter that can be increased to potentially improve designs, though it may also increase computational demands. Further, only a single post-PKS modification step was used for two reasons. First, to confirm that chimeric PKSs designed by RetroTide can reach the desired number of carbon atoms in commodity chemicals (from as little as 3 in acrylic acid to as many as 19 nonadecene) from just simple acyl-CoA building blocks, and without having to provide carbon atoms through monofunctional enzymes. Second, from an experimental perspective, fewer post-PKS modification steps minimize the likelihood of intermediate loss: while the enzymatic domains of type I PKSs are covalently connected and the growing carbon chain substrate is always tethered to a given module’s acyl carrier protein (ACP) domain through the ACP’s phosphopantetheinyl arm^25^, regular monofunctional enzymes do not enjoy such physical proximity in the cell^50,100^. In fact, it is common for engineered metabolic pathways to become disrupted when their intermediates are siphoned off into other pathways valuable for primary metabolism, ultimately resulting in low final product titres^50–57^. If successfully engineered, chimeric PKSs are less susceptible to such derailments. Recently, the unfolded protein linkers that covalently bind the catalytic domains of type I PKSs together have even been used to fuse together monofunctional enzymes in engineered metabolic pathways, leading to more than double the product titre in some cases^101^. Given such high substrate concentrations localized around individual catalytic domains rather than globally dispersed around the cell, PKSs also have the potential to force through reactions that may be thermodynamically uphill and would otherwise remain uncatalyzed by regular, monofunctional enzymes.

**Fig. 5:**
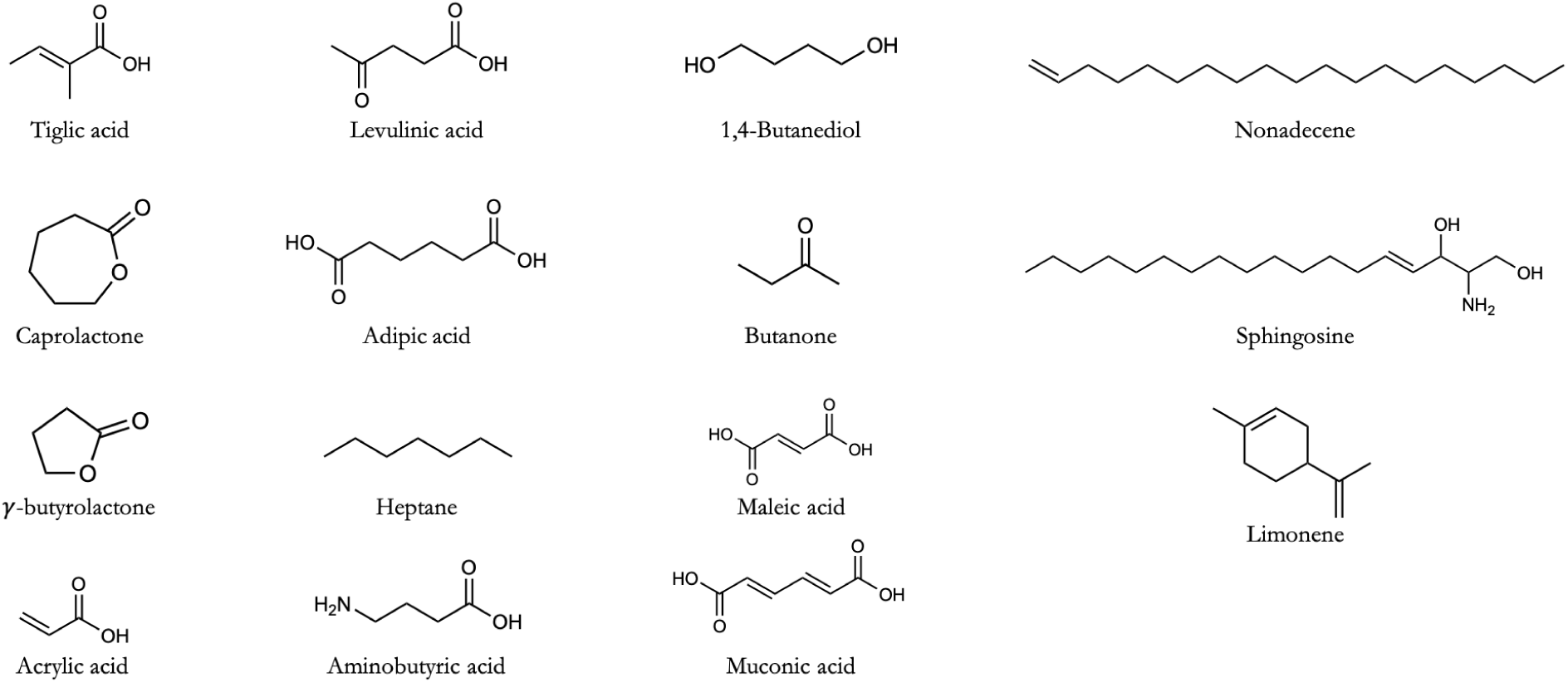
A structurally diverse set of 15 molecules were curated for the prototyping of BioPKS Pipeline. Fifteen molecules of varying carbon chain lengths and complexity were chosen as an initial prototyping set for BioPKS Pipeline to investigate if chimeric PKSs suggested by RetroTide would be able to construct the carbon backbones for each of these molecules. Both cyclic and aliphatic molecules were included in our set and amongst the aliphatic molecules, carbon backbones with as little as 3 (acrylic acid) and as many as 19 (nonadecene) carbon atoms were chosen to test the versatility of BioPKS Pipeline in constructing scaffolds of different lengths.

RetroTide was able to synthesize a carbon scaffold with at least as many carbon atoms as the final target for all 15 targets. Interestingly, for tiglic acid, caprolactone, and γ-butyrolactone, the top chimeric PKS designs were not only able to build the carbon backbones of these products but also accomplish the structural decorations necessary to synthesize all three targets completely (Fig. 6, SI Fig. 2). For the remaining 12 molecules, since PKSs did not synthesize the final target, a single-step post-PKS modification reaction was used with DORAnet. Amongst these 12 molecules, for acrylic acid, muconic acid, levulinic acid, adipic acid, 1,4-butanediol, and maleic acid, the top-PKS product serves as an excellent precursor for post-PKS modifications and these targets can be synthesized completely in a single post-PKS tailoring step (Fig. 6). For butanone, heptane, and nonadecene, however, the top PKS product does not lead to the target molecule after a single post-PKS modification step. Surprisingly, in these cases, a lower-ranked PKS design leading to a more structurally dissimilar intermediate actually serves as a better launchpad for the post-PKS modifications required to transform the corresponding PKS products into the final targets (Fig. 6). Here, we calculate chemical similarity by first computing the maximum common substructure (MCS) between a given (either PKS or post-PKS) product and the target chemical and then calculating the ratio of atoms in the MCS to the total number of unique atoms between both molecules, e.g. the MCS Tanimoto similarity. Finally, for limonene, aminobutyric acid, and sphingosine, BioPKS Pipeline was not able to reach these targets across all 6 PKS designs considered with a single monofunctional enzyme step. PKS designs for all successful and unsuccessful syntheses mentioned here can be found in SI Figs. 2-9.

**Fig. 6:**
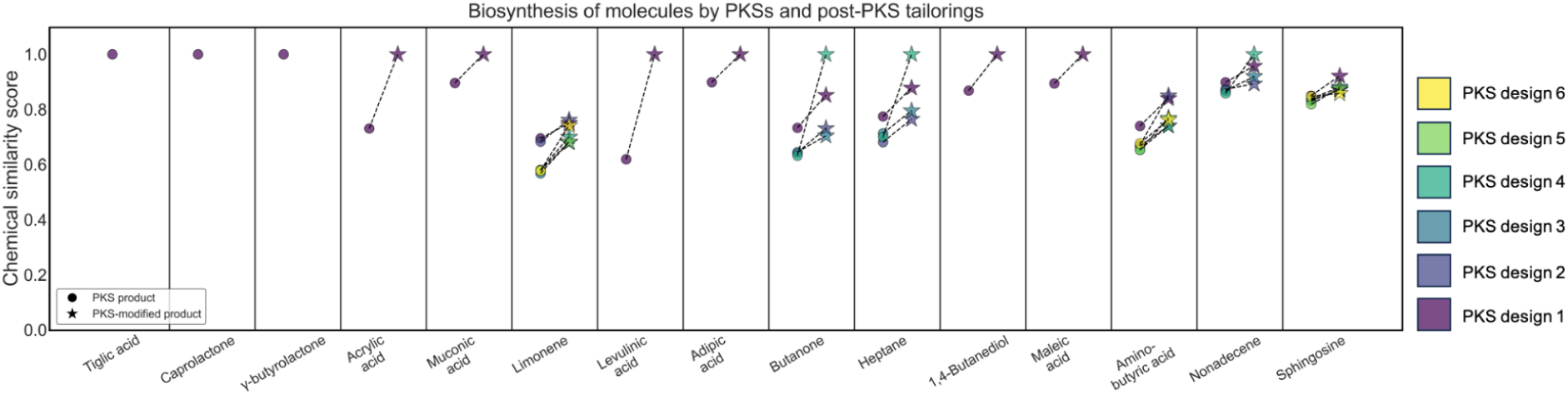
BioPKS Pipeline successfully constructs the carbon backbones of all 15 curated molecules and synthesizes 12 molecules completely. For 3 molecules out of the total 15 in our curated prototyping set (i.e. tiglic acid, caprolactone, γ-butyrolactone), BioPKS Pipeline synthesizes these molecules completely with the top-ranked type I polyketide synthase (PKS) design and does not require any further post-PKS modifications. For 9 molecules (acrylic acid, muconic acid, levulinic acid, adipic acid, butanone, heptane, 1,4-butanediol, maleic acid, and nonadecene), BioPKS Pipeline synthesizes these molecules completely using a single post-PKS modification step. Out of these 9 molecules, acrylic acid, muconic acid, levulinic acid, adipic acid, 1,4-butanediol, and maleic acid can be reached by a single-step modification of the top-ranked PKS design’s product while butanone, heptane, and nonadecene can only be reached by starting with a PKS product that is more structurally dissimilar with respect to the target molecule than the top-ranked PKS product. There exist 3 molecules out of the 15 total (limonene, aminobutyric acid, and sphingosine) that cannot be synthesized at all by BioPKS Pipeline within just one post-PKS step even after considering five alternate PKS designs.

### Validating BioPKS Pipeline against a large set of valuable biomanufacturing targets

In order to validate BioPKS Pipeline against a larger, unbiased collection of realistic biomanufacturing targets, and investigate the effect on target diversity of allowing extra post-PKS steps, we turned to a previously published set of 209 chemicals to be potentially produced via biomanufacturing^13^. Of these 209 chemicals, we filtered out C1 metabolites, cofactors, charged targets as well as any chlorinated or sulfonated species so as to yield a final validation set of 155 molecules. Cofactors were removed as target molecules because these would be consumed by DORAnet and C1 metabolites were also removed because their carbon backbone is too short for a PKS. Our reaction rules do not work on charged ions so these were removed as well. Of these 155 targets, 94 were non-aromatics while 61 were aromatics. We then prompted BioPKS Pipeline to synthesize all 155 chemicals with up to 2 post-PKS modification steps. For the 94 non-aromatic targets, we constrained BioPKS Pipeline to only using the aliphatic units of malonyl-CoA, methylmalonyl-CoA, hydroxyacetyl-CoA, methoxymalonyl-CoA, and allylmalonyl-CoA as starters and extenders, while for the 61 aromatic targets we allowed for all starters and extenders to be used. Note that not all of these substrates are naturally occurring in the positions used (e.g. as starters or extenders), see the section “Selection of starter and extender acyl-CoA derivatives by RetroTide” for our rationale on including them. In attempting to synthesize these molecules, stereochemical information was not considered and 5 alternate PKS designs beyond the top-ranked PKS design were considered.

BioPKS Pipeline successfully proposed designs to synthesize 93 molecules out of our total of 155, translating to a success rate of 60% (Fig. 7A). Of these 93 molecules, chimeric PKS-only designs from RetroTide could synthesize only 3 molecules exactly (Fig. 7A). Still, the carbon scaffolds of PKS products proved to be very valuable precursors for post-PKS tailoring. Within a single generation, DORAnet was able to synthesize 46 more molecules completely by building off of RetroTide’s carbon backbones. With a second generation, DORAnet was able to synthesize 44 more molecules (Fig. 7A), thereby reaching 93 molecules in total. For the remaining 62 molecules that could not be synthesized by BioPKS Pipeline, all top-ranked PKS products still comprised at least as many carbon atoms as their corresponding final targets. Moreover, for these 62 molecules, post-PKS modifications performed on the top-PKS product are still shown to improve the structural similarity of the post-PKS product over the PKS product when both are compared to their corresponding target molecule (Fig. 7B).

**Fig. 7:**
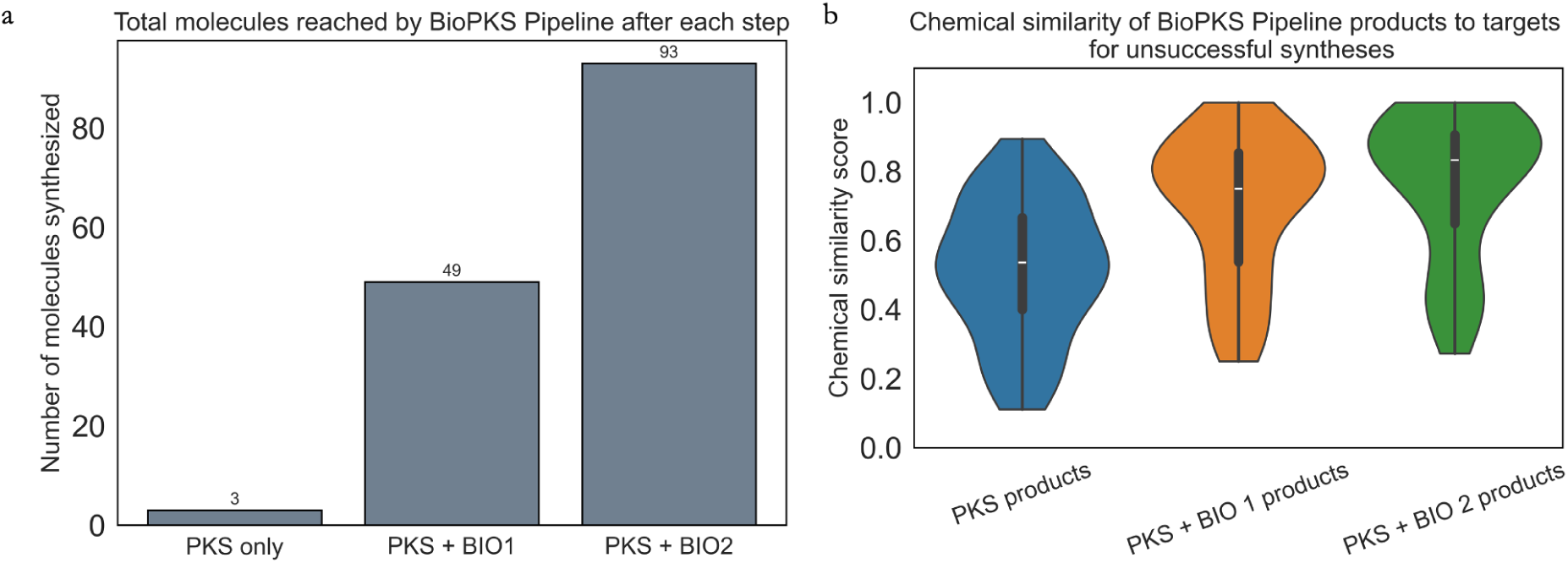
BioPKS Pipeline successfully suggests integrated biosynthetic pathways to 93 biomanufacturing candidates. **a** When BioPKS Pipeline is prompted to synthesize the validation set of 155 molecules that would be potential candidates for biomanufacturing, it achieves an overall hit-rate of 60% (i.e. 93/ 155). Of the 93 molecules that can be synthesized exactly, 3 are synthesized solely by PKSs while 46 more molecules can be synthesized with just one post-PKS tailoring step from the PKS product. After a second post-PKS step, 44 more molecules can be synthesized, bringing the total number of molecules synthesized by BioPKS Pipeline to 93. **b** Although the remaining 62 molecules out of 155 cannot be synthesized by BioPKS Pipeline, post-PKS modification steps are shown to improve the chemical similarity between the PKS product and the post-PKS product with respect to the target for these 62 target molecules that could not be synthesized.

We explored the possibility of attempting more than 2 post-PKS modifications on the PKS products involved in the synthesis of these remaining 62 molecules but ultimately decided against it. This is primarily because the increase in chemical similarity achieved with respect to the target is only marginal between the first and second post-PKS reactions as opposed to that between the PKS product and the first post-PKS modification step. This suggests that there are diminishing returns to increases in chemical similarity with each post-PKS step and that these 62 molecules may lie even farther outside the chemical space accessible by BioPKS Pipeline. Moreover, given that other known biological, chemical, or hybrid pathways to some of these molecules may already exist, if two post-PKS modification steps are not sufficient to synthesize these targets, then users may be able to find a more efficient pathway that does not rely on PKSs. Nonetheless, if users still want to use more than two post-PKS steps, perhaps for targets without any known pathways, then they can easily do so with the various pruning filters available in DORAnet. Since reaction networks grow exponentially with each generation, DORAnet enables users to discard intermediate metabolites that may appear too structurally dissimilar to the target, have more atoms than some predetermined maximum, or that simply fail to satisfy some other user-defined criteria. We emphasize here that users have the flexibility to easily build their own custom filters into DORAnet and prune pathways according to their own heuristics of interest. Ultimately, users can set up these filters to build post-PKS synthesis trees that are as deep or as shallow as they require.

### BioPKS Pipeline suggests novel pathways to molecules for which no known pathways exist

As a final stress-test on the robustness of our software, we prompted BioPKS Pipeline to suggest pathways to natural products for which no known pathways exist. We first attempted this for the antifungal cryptofolione isolated from the plant *Cryptocarya latifolia*^83^. The biosynthetic origin of cryptofolione has not been elucidated and only total synthesis pathways relying on synthetic chemistry have thus far been utilized to synthesize cryptofolione^102^. Given its therapeutic potential, cryptofolione represents a promising target for biomanufacturing with our ‘PKS-first’ approach presented here. When the structure of cryptofolione is input into BioPKS Pipeline without any stereochemical information, RetroTide suggested a PKS design consisting of 6 modules that could synthesize cryptofolione exactly (Fig. 8). This predicted chimera loads an aromatic starter unit, namely cinnamoyl-CoA to account for the aromatic moiety on one end of cryptofolione’s backbone. While not a known Type I PKS starter, cinnamoyl-CoA is a known type III PKS starter^30^, and it may be possible to engineer a Type I PKS that can accept it. The extension modules of this chimera then use only malonyl-CoA units to build the remainder of cryptofolione’s backbone. A terminal thioesterase (TE) domain that catalyzes an intramolecular cyclization reaction to form the lactone ring at the other end of cryptofolione’s backbone is finally used to complete cryptofolione’s proposed biosynthesis. Although the proposed cinnamoyl-CoA loading has not been observed in a type I PKS before, since the proposed extension modules here all use the commonly observed malonyl-CoA, we decided to search the ClusterCAD 2.0 database using these extension modules as queries. The goal of sending RetroTide’s suggested chimera for cryptofolione through ClusterCAD 2.0 was to see if existing PKSs could be found that are similar in domain architecture to the suggested chimera and whose parts could therefore be used to assemble the chimera. Our search revealed that module 1 of the simocyclinone PKS^103^ (MIBiG accession: BGC0001072.1) follows a KS-AT-DH-KR domain architecture and uses malonyl-CoA as an extender unit. This could be used to build modules 3 and 5 for the proposed chimeric PKS (Fig. 8). In a similar vein, module 2 of the simocyclinone PKS follows a KS-AT-KR domain architecture and could be used to build modules 1, 2, and 4 of the proposed PKS (Fig. 8). Since malonyl-CoA is a common extension unit, modules from other PKSs, such as the Niddamycin PKS^104^ (MIBiG accession: BGC0000113.1) can also be used to construct the suggested chimera.

**Fig. 8:**
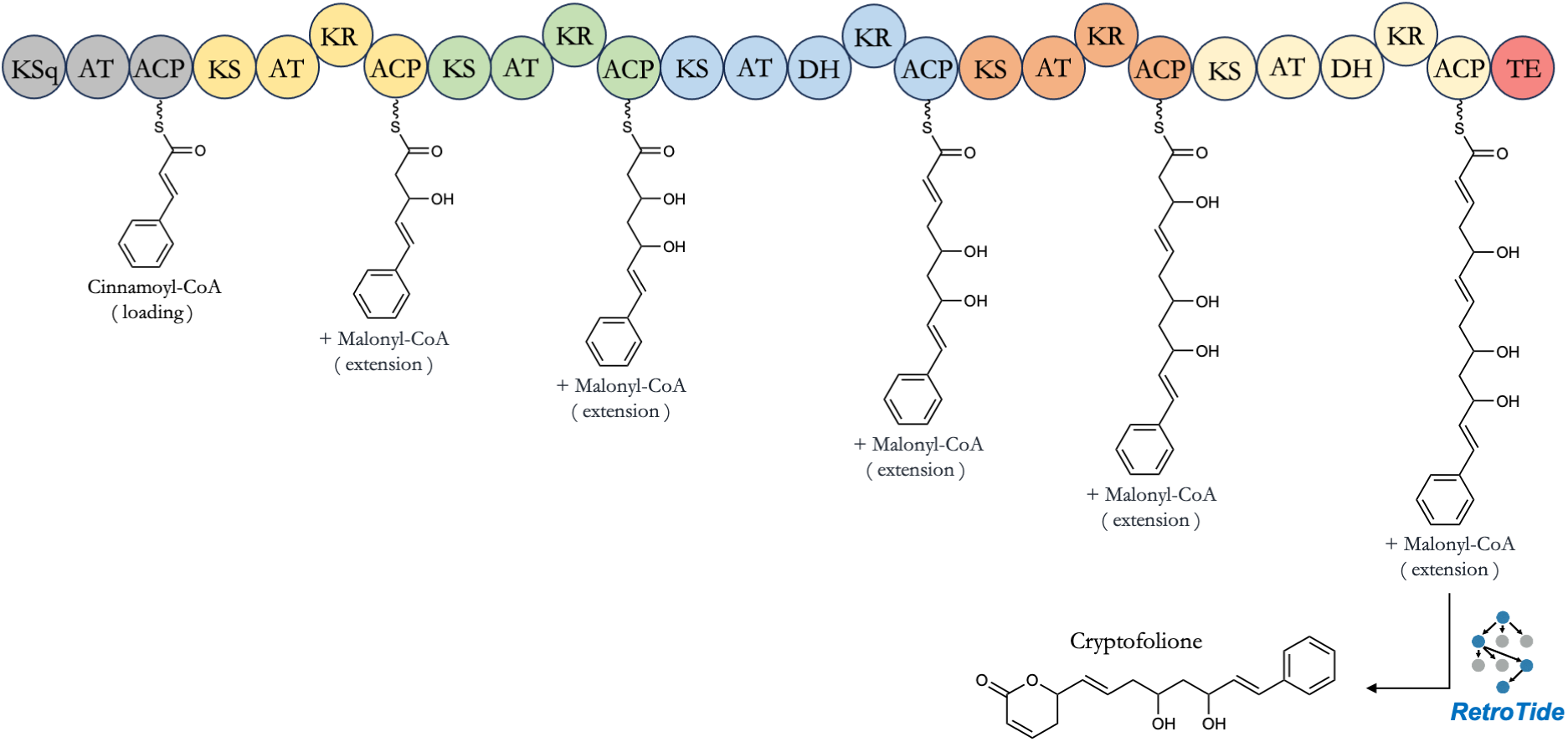
Proposed biosynthesis of cryptofolione by BioPKS Pipeline. When prompted to suggest a biosynthesis route to the natural product cryptofolione, a potential therapeutic whose biosynthetic origin remains unknown, BioPKS Pipeline calls upon RetroTide to successfully suggest a 6-module chimeric PKS that could synthesize cryptofolione exactly. This chimeric PKS selects cinnamoyl-CoA as a starter unit given the aromatic ring on one end of cryptofolione’s carbon backbone and subsequently only uses malonyl-CoA as extender units. The offloading reaction for the termination domain of this chimeric PKS is set to an intramolecular cyclization reaction so that the lactone ring on the other end of cryptofolione’s carbon backbone can be synthesized. Our PKS design lends insight into the possible biosynthetic origins of α-pyrones such as cryptofolione.

Since BioPKS Pipeline suggests a PKS design that can synthesize cryptofolione exactly, no post-PKS modification steps are needed. We note that in our proposed synthesis of cryptofolione, the alkene group adjacent to the acyl group prior to offloading is in the trans configuration while after lactonization, it changes to the cis configuration. We are uncertain if thioesterase domains that perform cyclizations would be able to invert double bond stereochemistry as, to our knowledge, no such examples have yet been observed in nature. While in this work, we focused on harnessing PKSs and enzymatic reactions to obtain the desired two-dimensional target structure, future work will incorporate more stereochemistry corrections such that even the desired three-dimensional structure is produced.

We then prompted BioPKS Pipeline to suggest a synthesis for the natural product basidalin. Basidalin is a smaller but potent antibiotic first extracted in 1983 from the fungus *Leucoagaricus naucina*^84^. Despite its early extraction, the first total chemical synthesis of basidalin was only achieved recently in 2016^105^. As is the case with cryptofolione, the biosynthetic origins of basidalin have yet to be uncovered. Thus, we prompted BioPKS Pipeline to synthesize basidalin and the suggested PKS design was able to synthesize basidalin’s carbon backbone exactly (Fig. 9). This predicted chimera features 3 modules in total. The loading module and first extension module both utilize hydroxymalonyl-CoA as starter and extender units respectively while the final extension module utilizes malonyl-CoA as an extender unit. Similar to the predicted chimera for the earlier proposed biosynthesis of cryptofolione, a TE domain that catalyzes an intramolecular cyclization reaction is used for the proposed biosynthesis of basidalin. Since basidalin is a high-value molecule without any known biosynthetic pathways, we decided to use three post-PKS modification steps in its synthesis and were ultimately able to synthesize basidalin exactly. The first suggested post-PKS reaction features a peroxidase, the second features an aminotransferase, and the third involves an alcohol dehydrogenase. Interestingly, while the first and second post-PKS reactions by DORAnet have a high predicted feasibility score and are thermodynamically downhill, the last post-PKS reaction is predicted to be very infeasible and found to be thermodynamically uphill. Since all reaction free energies are calculated under typical cellular conditions of 298K, ionic strength 0.25M, pH 7.4, and pMg 3.0 for all species at 1 M concentrations (see Methods), users may want to perform such uphill reactions *in vitro* wherein these conditions may be tweaked a little further to improve thermodynamic favorability. The ability of BioPKS Pipeline to suggest pathways to such complex natural products could also enable users to utilize BioPKS Pipeline to annotate orphan biosynthetic gene clusters.

**Fig. 9:**
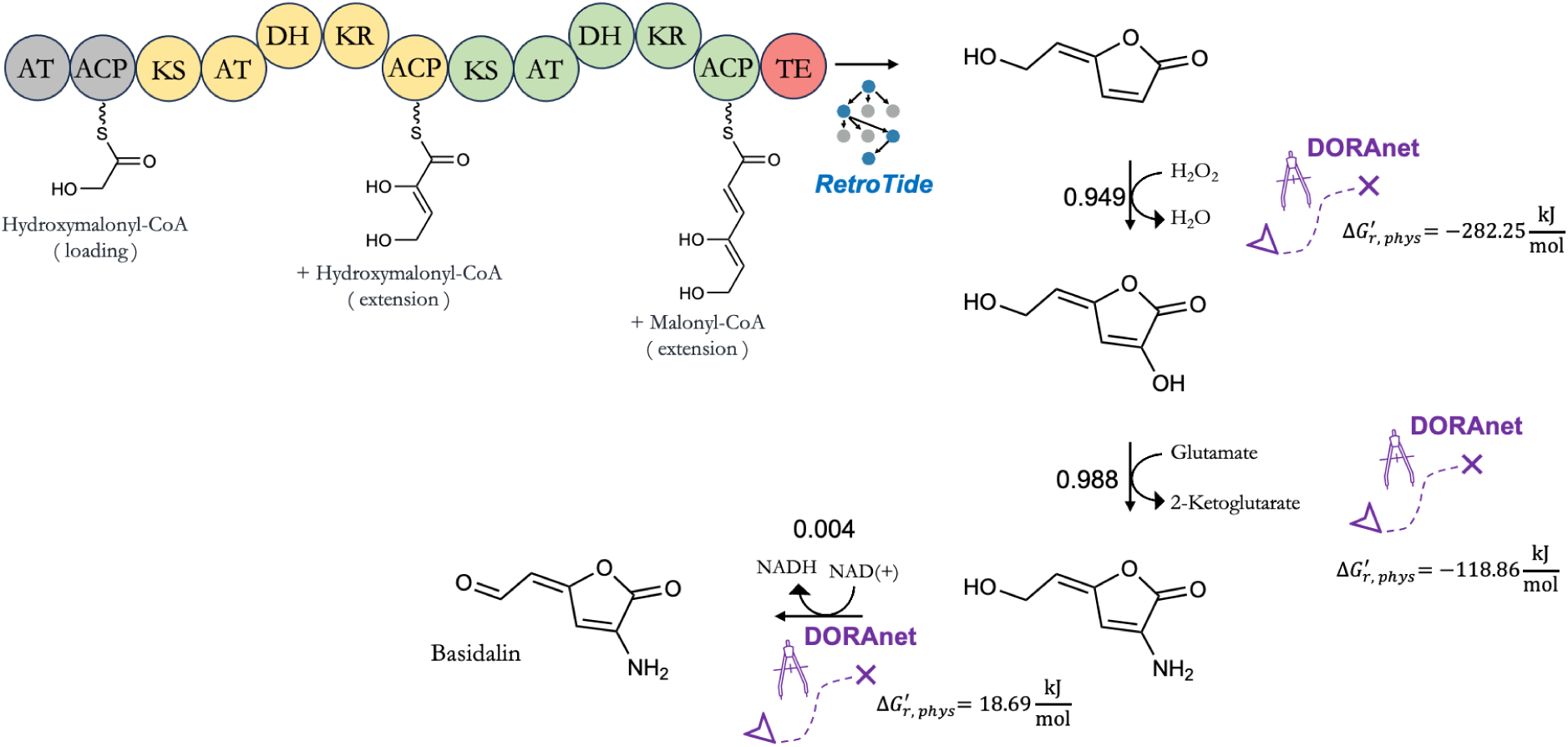
Proposed biosynthesis of Basidalin by BioPKS Pipeline. When prompted to suggest a biosynthesis route to basidalin, another therapeutic natural product whose biosynthetic origin also remains unknown, BioPKS Pipeline suggests a chimeric PKS design and three post-PKS modification steps to synthesize basidalin exactly. This chimeric PKS consists of 3 modules. The loading module and first extension module both use hydroxymalonyl-CoA as starter and extender units respectively while the final extension module uses malonyl-CoA as an extender unit. Again, an intramolecular cyclization reaction is chosen for offloading and we note that RetroTide is able to synthesize the γ-lactone backbone of basidalin exactly. Subsequently, three post-PKS modification steps involving peroxygenase and aminotransferase enzymes can produce the target basidalin product.

## Discussion

Here, we present a retrobiosynthesis software that harnesses both multifunctional and monofunctional enzymes to propose the *de novo* biosynthesis of a broad spectrum of chemical targets - from simple commodity chemicals to more complex natural products for which biosynthetic pathways may or may not already exist. Crucially, we have also put forth a valuable paradigm for the synthesis of such molecules - that multifunctional enzymes, such as type I PKSs, can be exploited for creating the carbon scaffolds of target products while regular, monofunctional enzymes within biology can subsequently be brought in for structural refinements that eventually transform such scaffolds into the target product exactly.

Our motivation for developing this ‘PKS-first’ algorithm to drive BioPKS Pipeline lies in expanding biologists’ capabilities to explore chemical space by complimenting the strengths of PKSs with that of monofunctional enzymes. PKSs excel at synthesizing elongated carbon backbones and/ or incorporating unique functional handles into target molecules from a common pool of simple, acyl-CoA building blocks. Monofunctional enzymes, by contrast, rarely catalyze such iterative C-C bond formation reactions but are adept at precise, regioselective functional group transformations. Indeed, there are several reaction chemistries, such as glycosylations, monooxygenations, and transaminations that PKSs are simply unable to perform and must instead rely on monofunctional enzymes. Rather than attempting to manufacture chemicals using only one of these routes, we propose that a wider space of molecules can be accessed by integrating the two.

In this study, we demonstrated the effectiveness of our approach and of BioPKS Pipeline across various test-cases of increasing complexity. First, we showcased BioPKS Pipeline’s two modes of operation by synthesizing 4-hydroxybutyric acid completely through PKSs, while 2-pentanone was reached with just one post-PKS modification step. Although metabolic pathways have already been engineered for the biosynthesis of both chemicals, our proposed pathways offer a compelling alternative that may minimize intermediate loss, a common occurrence with classic metabolic pathways. Subsequently, we further prototyped BioPKS Pipeline for the synthesis of 15 structurally diverse chemicals and proved that, for certain targets, considering alternate PKS designs beyond the top-ranked design can yield better intermediates for post-PKS modifications to synthesize the final target. After these initial tests, we turned to a published set of molecular candidates for biomanufacturing and proved that we could synthesize 60% of them (93 out of 155) with our pipeline while allowing for up to two post-PKS modification steps. BioPKS Pipeline can access an even larger chemical space by considering further post-PKS steps. Longer pathways, however, will certainly be more challenging to successfully build experimentally, so we decided to limit our analysis to two post-PKS steps so that our proposed designs are of a realistic complexity. As a final test of BioPKS Pipeline, we prompted it to suggest combined PKS and post-PKS pathways to complex natural therapeutics for which biosynthetic gene clusters have not been elucidated. In doing so, BioPKS Pipeline was able to suggest novel pathways to the natural products cryptofolione (antifungal) and basidalin (antibiotic) for which no known biosynthetic pathways exist.

One limitation of our tool is the lack of support for stereochemistry corrections. While the enantioselective synthesis of compounds is a key strength of PKSs^106,107^ and many monofunctional enzymes^108^, a number of retrobiosynthesis tools in the literature cannot reliably predict the correct stereoisomer of both existing and novel reactions. Nonetheless, with the recent development of cheminformatics tools capable of both extracting and applying stereochemistry-aware reaction templates^109–111^, we expect our future work to address such corrections as well.

Overall, while the successful engineering and construction of chimeric PKSs remains an ongoing challenge, our *in silico* pipeline highlights the potential of the field, given the diversity of products that can be synthesized by integrating PKSs with post-PKS pathways. In addition to suggesting pathway chemistries to target molecules using both PKSs and monofunctional enzymes, BioPKS Pipeline also enables users to rank results through reaction thermodynamics and feasibility calculations to observe if potentially promiscuous, wild-type enzymes would catalyze predicted reactions or if such enzymes would need to be mutated further. With all of these functionalities built in, BioPKS Pipeline serves as a flexible and comprehensive tool for users looking to synthesize molecules via both PKSs and monofunctional enzymes. We have also significantly modularized the code for BioPKS Pipeline such that if users wish, other tools in the literature could be further bundled in for various post-processing analyses involving enzyme selection^112–115^, global cofactor usage in the cell^116^, and predicting potential enzyme cross-talk^117^. Moreover, with the abundance of enzyme engineering techniques now available - both experimental^118,119^ and computational^120–123^ - our designs here serve as a valuable starting point for the research community to continue iterating on them. For users who may be interested, we have also provided several tutorials on our Github repository that show more examples of using BioPKS Pipeline in various scenarios.

## Methods

### User-defined inputs for BioPKS Pipeline

Users can initialize a BioPKS Pipeline object in the Python programming language with the simplified molecular input linear entry system (SMILES) string of their target chemical (Fig. 1). Users can also specify if this target chemical should be synthesized using either only PKSs, or only monofunctional enzymes, or both. In order to generate PKS designs, users can either allow for all 30 starter (SI Table 1) and 11 extender acyl-CoA (SI Table 2) units available within BioPKS Pipeline to be used as building blocks or choose a specific subset of these units. BioPKS Pipeline supports both an intramolecular cyclization as well as a thiolysis reaction to release the bound PKS product from the designed PKS’s thioesterase (TE) domain and users can select from either of these offloading reactions to generate their PKS product. When using combined PKS and non-PKS pathways, users must specify the number of post-PKS modification steps to perform on the PKS product.

Currently, BioPKS Pipeline does not support any stereochemistry corrections on-the-fly. This is because there are still only a few fingerprinting methods and chemical similarity metrics that can accurately compare the three-dimensional structural similarity of two chiral molecules. Further, since different chemical similarity metrics emphasize different features within a molecule anyway, in this work, we prioritized obtaining the correct two-dimensional structure of the target molecule. Consequently, all molecules presented here are shown without their stereochemical configurations. With recent work on chiral fingerprints published in the literature^124^, however, we expect our future work to at least be able to implement stereochemistry corrections in a post-hoc fashion.

We emphasize here that BioPKS Pipeline does not currently support invoking PKSs a second time if a target molecule cannot be reached using PKSs first and monofunctional enzymes second. While such an approach that relies on PKSs interchangeably would almost certainly expand the net chemical space accessible, this would be too difficult to implement experimentally within the laboratory since PKSs only accept a very limited pool of substrates. Consequently, it may not be biologically practical to build a scaffold with a PKS, modify it enzymatically, and then convert it into an acyl-CoA derivative for use by a PKS.

### Computational design of chimeric type I PKSs with RetroTide

Chimeric type I PKS designs are created using RetroTide, the first key component within BioPKS Pipeline. Upon providing the simplified molecular input linear entry system (SMILES) string of a target molecule, RetroTide first attempts to design PKSs that can synthesize this molecule using various modules, domain architectures, starter units, and extender units (Fig. 10). Users can customize which starter and extender units they would like to use in building chimeric PKSs from a provided list of 30 starter (SI Table 1) and 11 extender (SI Table 2) units, or optionally expand the list themselves. The complete list of starter and extender units available can be found in the SI. RetroTide builds PKS designs by recursively adding modules, thereby synthesizing in the forward direction (Fig. 9). Within each round of adding a module, the PKS product is computed and only the top N designs leading to the most chemically similar products progress onto the next recursive call. Here, N is a hyperparameter that can be set by the user and has a default value of 15. The specific metric of chemical similarity used by RetroTide to compute the structural similarity between the current PKS product and a target product is also an option that can be selected by the user. Users can choose between either computing the (1) Atom-Atom-Path similarity^125^, the (2) Atom pair Tanimoto similarity^126^, or the (4) Tanimoto similarity using the maximum common substructure between the current PKS product and the target product either with or without chirality. After RetroTide has completely finished designing chimeric PKSs, these designs are returned in descending order of chemical similarity - as measured between each design’s corresponding product and the target product. Users can then decide between terminating the growing carbon chain on the simulated chimeric PKS via either an intramolecular cyclization reaction, such as that involved in the formation of 6-deoxyerythronolide B^25^, or a thioesterase reaction, thereby forming a terminal carboxylic acid group. The highest-ranked PKS design from RetroTide either may or may not have produced the user’s target product, depending on whether this product is reachable by purely PKS chemistry.

**Fig. 10:**
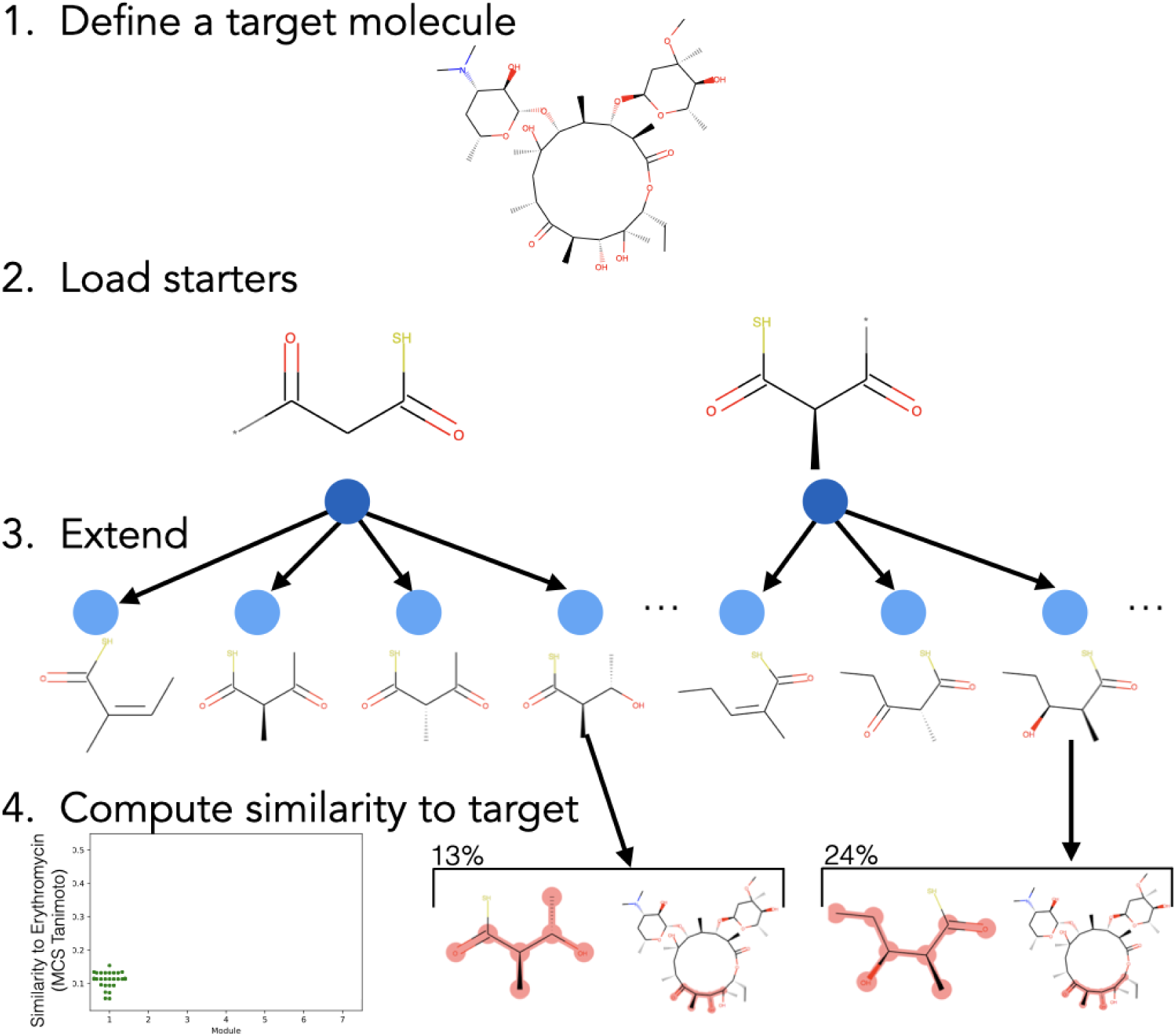
RetroTide algorithm for the design of PKSs from a target chemical structure. The RetroTide algorithm has the following steps: **1)** The user provides a desired target chemical structure. **2)** RetroTide loads all possible PKS starter module final structures. **3)** The cartesian product between all possible starters, and all possible extension modules is used to compute all possible 1 module PKS products. These extension module structures are pre-computed once and reused as RetroTide runs. **4)** All structures are compared with the user defined similarity metric to the defined target, and only the N (a user selectable parameter) top scoring designs are retained. **5)** Steps 2-4 are repeated iteratively adding all possible single PKS modules to the end of each of the best scoring designs from the previous round, until scores stop increasing for each round and then the algorithm halts, providing the best scoring designs to the user. **6)** The best scoring designs are optionally ‘released’ as lactone/lactam, or carboxylic acid products.

With the PKS designs obtained from RetroTide, users can search the ClusterCAD 2.0 PKS database^85^ (freely available online at https://clustercad.jbei.org/pks/) for naturally-occurring designs that match closely with the suggested chimeric designs, so as to select an initial design for biological engineering. We also note RetroTide is cis/trans agnostic and since recent work has shown that trans-ATs can be engineered^127^, we encourage users to implement RetroTide designs as trans-AT PKSs and/or supplement the list of PKS starters and extenders with additional substrates, as appropriate for a given biosynthesis goal.

### Performing post-PKS modifications on RetroTide products with DORAnet

Post-PKS modifications are performed using our DORAnet platform available at https://github.com/wsprague-nu/doranet. DORAnet is built on the concept of our previously released metabolic network expansion tool, Pickaxe^128^. If the user-defined target product was not reached by RetroTide, the PKS product is funneled as a starting molecule into DORAnet. DORAnet is a retrosynthesis tool designed to utilize both catalytic and metabolic reaction rules for discovering and ranking hybrid pathways. Further details about DORAnet will be published in a separate work. In this work, DORAnet was used to generate metabolic *in-silico* network expansions (MINEs) by recursively expanding upon a precursor metabolite using reaction rules. Users can decide between two sets of reaction rules in generating MINEs. The first set of rules to choose from is our generalized and maximally promiscuous JN1224MIN rule set^86^, which predicts promiscuous reactions by only considering the substrate chemical moieties that can directly undergo enzymatic reactions. The second, is our intermediate JN3604IMT rule set^87^, which not only considers reacting moieties in predicting reactions but also the surrounding chemical neighborhoods within which these moieties are present. Both of these rule sets were generated from MetaCyc and approximately cover 85% of all chemistries encapsulated within BRENDA and KEGG. In performing post-PKS modifications, users must specify the number of post-PKS steps, i.e., the number of generations for which DORAnet would be run. After post-PKS modifications, if the target product is reached, pathways between the PKS product and the final, downstream target are returned. If the target product is not reached, however, then pathways to the most chemically similar non-PKS product with respect to the final target are returned. When running longer post-PKS modification pathways, users also have the option to choose from various in-built DORAnet compound filters (such as the molecular weight filter or the chemical similarity filter), so as to ensure that MINEs remain computationally tractable.

Once post-PKS pathways have been elucidated, users can use the specific reaction rules that were involved in generating predicted reactions to search for UniProt identification numbers^129^ of potentially promiscuous enzymes present within either JN1224MIN^86^ or JN3604IMT^87^, that may be able to catalyze predicted reactions.

### Machine-learning guided selection of post-PKS enzymatic pathways

Depending on the number of post-PKS modification steps performed, the number of post-PKS pathways found between the PKS product and the final target may be too large to conceivably manually analyze in order to elucidate the most feasible pathway chemistries for experimentation. As such, our previously released enzymatic reaction feasibility classifier^91^ (available at https://github.com/tyo-nu/DORA_XGB) has been bundled into BioPKS Pipeline to further rank post-PKS pathways in terms of their predicted feasibility scores (Fig. 2). Feasibility scores are computed for each reaction in a pathway by considering the SMILES strings of all participating species, i.e., substrates, products, and cofactors. Our reaction feasibility model is based on the gradient boosted trees architecture and was trained using reported thermodynamically feasible reactions as positive data and thermodynamically infeasible as well as synthetically generated infeasible reactions as negative data. This allows the computed feasibility score to reflect both the thermodynamic feasibility of a reaction as well as the likelihood of an enzyme transforming a certain moiety at a given reaction site. After computing the feasibility score for all reactions in a pathway, a net feasibility score for the entire pathway is computed by taking the product of all constituent reactions’ feasibility scores. Instead of using the feasibility score, which is a continuous number between 0 and 1, users can also opt for a binary label on whether a predicted reaction would be feasible or not using the built-in feasibility thresholds for the feasibility classifier. We note here that even though we have used our DORA-XGB model to rank DORAnet reactions and pathways, users are able to use other metrics as well for pathway ranking. Beyond reaction feasibility and thermodynamics, this can even include reaction/ pathway novelty, or pathway length. In future work, a multi-objective optimization involving all of these metrics using a different search algorithm, such as A* or monte carlo tree search may help to expand the accessible chemical space even further.

### Thermodynamic calculations of post-PKS enzymatic pathways

In order to further down-select predicted pathways, BioPKS Pipeline also enables users to calculate the Gibbs free energies of reactions as well as the maximum/minimum driving force (MDF) of multi-step pathways^130^. The open-source software eQuilibrator 3.0^81^ was used to calculate the change in Gibbs free energy due to a reaction, Δ_*r*_*G*’. eQuilibrator 3.0 uses the component contribution method for first estimating the standard Gibbs free energy change due to a reaction, Δ_*r*_*G*°, by assuming reactant concentrations of 1 M. These Δ_*r*_*G*° values are then transformed to our prespecified conditions of temperature 298 K, ionic strength 0.25 M, pH 7.4, and pMg 3.0 to calculate Δ_*r*_*G*’ under common cellular conditions. Users can also adjust these thermodynamic parameters in their calculations as desired. In order to ensure that BioPKS Pipeline is accessible to all, we did not incorporate eQuilibrator-assets into our software and only incorporated the general eQuilibrator API. Thus, if there exists a new compound not present within the eQuilibrator compound database, it cannot be decomposed into its groups and added to the database linked below since doing so would require a ChemAxon license to calculate the pKa values of new compounds. Consequently, the change in Gibbs free energy due to a reaction, Δ_*r*_*G*’ can only be computed for predicted DORAnet reactions in which all participating compounds are present in the eQuilibrator compound database, which in turn can be obtained as a SQLite database through Zenodo (zenodo.org/records/4128543).

Provided that the change in Gibbs free energy due to a reaction, Δ_*r*_*G*’ can be computed for all reactions along a multi-step pathway, users can subsequently calculate the MDF of a post-PKS pathway. This calculation seeks a set of metabolite concentrations that minimizes the Δ_*r*_*G*’ value of the most thermodynamically uphill or bottlenecked reaction within a multi-step pathway.

### Mining of potential bio-based chemicals set from a previously published set

In order to collect a larger set of chemicals upon which we could validate BioPKS Pipeline, we extracted 198 chemicals from a list of 209 chemicals that could hypothetically be produced by biomanufacturing as provided by Wu and colleagues through genome scale models of *Saccharomyces cerevisiae* and *Escherichia coli*^13^. Not all 198 chemicals extracted have existing engineered metabolic pathways for their production. Of these 198 chemicals, we filtered out C1 metabolites, common cofactors, and any sulfonated or chlorinated species, leaving us with 155 molecules in total. With these remaining targets, we queried their KEGG identifiers against the KEGG database^89^ using their application programming interface (API) and were able to receive SMILES strings for all 155 molecules. For these 155 molecules, post-PKS modifications of up to two steps were used and pathways were identified by comparing the SMILES strings of post-PKS pathways exactly with that of targets.

## Code Availability

BioPKS Pipeline and RetroTide are provided as free open source software at https://github.com/JBEI/BioPKS-Pipeline and https://github.com/JBEI/RetroTide respectively. All notebooks used in generating figures presented here can be found on the BioPKS Pipeline github repository at: https://github.com/JBEI/BioPKS-Pipeline/tree/main/notebooks.

## Data Availability

All data used here can be found on the BioPKS Pipeline github repository at: https://github.com/JBEI/BioPKS-Pipeline/tree/main/data.

## Acknowledgements

This work was funded in part by the DOE Joint BioEnergy Institute, supported by the U. S. Department of Energy Office of Science’s Biological and Environmental Research (BER) program, through contract DE-AC02-05CH11231 between Lawrence Berkeley National Laboratory and the U.S. Department of Energy. Funding for Yash Chainani for this study was partly provided by the Northwestern University Graduate School Cluster in Biotechnology, Systems, and Synthetic Biology, which is affiliated with the Biotechnology training program. The funding for Margaret Guilarte-Silva was partly provided by the Northwestern University DeFeo Family Undergraduate Research Fellowship in Synthetic Biology and partly by the Northwestern University Summer Undergraduate Research Award. The authors would like to acknowledge Matthew Incha and Patrick Kinnuen for their reading of the manuscript and corresponding feedback. This research project was supported in part through the computational resources and staff contributions provided by the Quest high performance computing facility at Northwestern University, which is jointly supported by the Office of the Provost, the Office of Research, and Northwestern University Information Technology. This research also used resources of the National Energy Research Scientific Computing Center (NERSC), a Department of Energy Office of Science User Facility using NERSC award ERCAP0028489.

## Author Contributions

**Yash Chainani:** Methodology and Software, Data curation, Writing – original draft and subsequent edits. Visualization – generated the figures in this manuscript. **Jacob Diaz:** Methodology and Software, **Margaret Guilarte-Silva:** Methodology and Software, **Vincent Blay:** Methodology and Software, **Quan Zhang:** Methodology and Software, **William Sprague:** Methodology and Software, **Keith E. J. Tyo:** Writing – review & editing, were the principal investigators who directed this project, contributed to the data analysis and interpretation, as well as edited the manuscript. **Linda J. Broadbelt:** Conceptualization – conceptualized the approach of this study. Investigation. Formal analysis. Writing – review & editing, were the principal investigators who directed this project, contributed to the data analysis and interpretation, as well as edited the manuscript. **Aindrila Mukhopadhyay:** Conceptualization – conceptualized the approach of this study. Investigation. Formal analysis. Writing – review & editing, were the principal investigators who directed this project, contributed to the data analysis and interpretation, as well as edited the manuscript. **Jay D. Keasling:** Conceptualization – conceptualized the approach of this study. Investigation. Formal analysis. Writing – review & editing, were the principal investigators who directed this project, contributed to the data analysis and interpretation, as well as edited the manuscript. **Hector Garcia Martin:** Conceptualization – conceptualized the approach of this study. Investigation. Formal analysis. Writing – review & editing, were the principal investigators who directed this project, contributed to the data analysis and interpretation, as well as edited the manuscript. **Tyler W. H. Backman:** Methodology and Software, Data curation, Conceptualization – conceptualized the approach of this study. Investigation. Formal analysis. Writing – review & editing, were the principal investigators who directed this project, contributed to the data analysis and interpretation, as well as edited the manuscript.

## References

1. Keasling, J. D. Manufacturing Molecules Through Metabolic Engineering. Science 330, 1355–1358 (2010).

2. Clomburg, J. M., Crumbley, A. M. & Gonzalez, R. Industrial biomanufacturing: The future of chemical production. Science 355, aag0804 (2017).

3. Choi, K. R., Jang, W., Yang, D., Cho, J. S., Park, D., Lee, S. Y. Systems metabolic engineering strategies: integrating systems and synthetic biology with metabolic engineering. Trends Biotechnol. 37, 817–837 (2019).

4. Ko, Y.-S., Kim, J. W., Lee, J. A., Han, T., Kim, G. B. et al. Tools and strategies of systems metabolic engineering for the development of microbial cell factories for chemical production. Chem. Soc. Rev. 49, 4615–4636 (2020).

5. Fackler, N., Heffernan, J. K., Boock, J. T., Gözmen, B., Schmitz, L. M., Köpke, M., & Jewett, M. C. Stepping on the gas to a circular economy: accelerating development of carbon-negative chemical production from gas fermentation. Annu. Rev. Chem. Biomol. Eng. 12, 439–470 (2021).

6. The White House. Executive order on advancing biotechnology and biomanufacturing innovation for a sustainable, safe, and secure American bioeconomy. The White House (2022).

7. Boston Consulting Group (BCG). Synthetic biology is about to disrupt your industry. BCG Global (2022). Available at: https://www.bcg.com/publications/2022/synthetic-biology-is-about-to-disrupt-your-industry.

8. Boston Consulting Group (BCG). Synthetic biology is getting closer to industrial scale. BCG Global (2024). Available at: https://www.bcg.com/capabilities/digital-technology-data/emerging-technologies/expert-insights/nicolas-goeldel.

9. Lawson, C. E., Martí, J. M., Radivojević, T., Jonnalagadda, S. V. R., Gentz, R. et al. Machine learning for metabolic engineering: a review. Metab. Eng. 63, 34–60 (2021).

10. Carbonell, P., Radivojevic, T. & García Martín, H. Opportunities at the Intersection of Synthetic Biology, Machine Learning, and Automation. ACS Synth. Biol. 8, 1474–1477 (2019).

11. Kumar, A., Wang, L., Ng, C. Y. & Maranas, C. D. Pathway design using de novo steps through uncharted biochemical spaces. Nat Commun 9, 184 (2018).

12. Maitra, S. & Maitra, K. Chemistry of bioproducts. In Practices and Perspectives in Sustainable Bioenergy: A Systems Thinking Approach (eds. Mitra, M. & Nagchaudhuri, A.) 233–267 (Springer India, New Delhi, 2020).

13. Wu, W., Long, M. R., Zhang, X., Reed, J. L. & Maravelias, C. T. A framework for the identification of promising bio-based chemicals. Biotechnol. Bioeng. 115, 2328–2340 (2018).

14. Yim, H., Haselbeck, R., Niu, W., Pujol-Baxley, C., Burgard, A. et al. Metabolic engineering of *Escherichia coli* for direct production of 1,4-butanediol. Nat. Chem. Biol. 7, 445–452 (2011).

15. Raab, A. M., Gebhardt, G., Bolotina, N., Weuster-Botz, D. & Lang, C. Metabolic engineering of *Saccharomyces cerevisiae* for the biotechnological production of succinic acid. Metab. Eng. 12, 518–525 (2010).

16. Lee, S. Y., Kim, H. U., Chae, T. U., Cho, J. S., Kim, J. W. et al. A comprehensive metabolic map for production of bio-based chemicals. Nat. Catal. 2, 18–33 (2019).

17. Jang, W. D., Kim, G. B. & Lee, S. Y. An interactive metabolic map of bio-based chemicals. Trends Biotechnol. 40, 1308–1311 (2022).

18. Matar, S. & Hatch, L. F. Chemistry of Petrochemical Processes. (Elsevier, 2001).

19. Sirirungruang, S., Markel, K. & Shih, P. M. Plant-based engineering for production of high-valued natural products. Nat. Prod. Rep. 39, 1492–1509 (2022).

20. Jensen, P. R. Natural products and the gene cluster revolution. Trends Microbiol. 24, 968–977 (2016).

21. Schläpfer, P., Zhang, P., Wang, C., Kim, T., Banf, M. et al. Genome-wide prediction of metabolic enzymes, pathways, and gene clusters in plants. Plant Physiol. 173, 2041–2059 (2017).

22. Rokas, A., E. Mead, M., L. Steenwyk, J., A. Raja, H. & H. Oberlies, N. Biosynthetic gene clusters and the evolution of fungal chemodiversity. Nat. Prod. Rep. 37, 868–878 (2020).

23. Grininger, M. Enzymology of assembly line synthesis by modular polyketide synthases. Nat. Chem. Biol. 19, 401–415 (2023).

24. Nivina, A., Yuet, K. P., Hsu, J. & Khosla, C. Evolution and diversity of assembly-line polyketide synthases: focus review. Chem. Rev. 119, 12524–12547 (2019).

25. Khosla, C. Structures and mechanisms of polyketide synthases. J. Org. Chem. 74, 6416–6420 (2009).

26. Khosla, C., Tang, Y., Chen, A. Y., Schnarr, N. A. & Cane, D. E. Structure and mechanism of the 6-deoxyerythronolide B synthase. Annu. Rev. Biochem. 76, 195–221 (2007).

27. Wagner, T. D., Trauner, D., Mihoreanu, L., Koert, U. & Hertweck, C. α-Methylation follows condensation in the gephyronic acid modular polyketide synthase. Chem. Commun. 52, 8822–8825 (2016).

28. Young, J., Trauner, D., Mikleušević, G., Mihoreanu, L., Koert, U. & Hertweck, C. Elucidation of gephyronic acid biosynthetic pathway revealed unexpected SAM-dependent methylations. J. Nat. Prod. 76, 2269–2276 (2013).

29. Hertweck, C., Luzhetskyy, A., Rebets, Y. & Bechthold, A. Type II polyketide synthases: gaining a deeper insight into enzymatic teamwork. Nat. Prod. Rep. 24, 162–190 (2007).

30. Yu, D., Xu, F., Zeng, J. & Zhan, J. Type III polyketide synthases in natural product biosynthesis. IUBMB Life 64, 285–295 (2012).

31. Keatinge-Clay, A. T. The structures of type I polyketide synthases. Nat. Prod. Rep. 29, 1050 (2012).

32. Li, H., Cann, A. F. & Liao, J. C. Biofuels: Biomolecular Engineering Fundamentals and Advances. Annu. Rev. Chem. Biomol. Eng. 1, 19–36 (2010).

33. Menzella, H. G., Reid, R., Carney, J. R., Chandran, S. S., Reisinger, S. J., Patel, K. G. & Santi, D. V. Combinatorial polyketide biosynthesis by de novo design and rearrangement of modular polyketide synthase genes. Nat. Biotechnol. 23, 1171–1176 (2005).

34. McDaniel, R., Ebert-Khosla, S., Hopwood, D. A. & Khosla, C. Engineering broader specificity into an antibiotic-producing polyketide synthase. Science 279, 199–202 (1998).

35. Zhu, X., Liu, J. & Zhang, W. *De novo* biosynthesis of terminal alkyne-labeled natural products. Nat Chem Biol 11, 115–120 (2015).

36. Porterfield, W. B., Poenateetai, N. & Zhang, W. Engineered biosynthesis of alkyne-tagged polyketides by type I PKSs. iScience 23, 100943 (2020).

37. Musiol-Kroll, E. M., Zubeil, F., Schafhauser, T., Härtner, T., Kulik, A. et al. Polyketide bioderivatization using the promiscuous acyltransferase KirCII. ACS Synth. Biol. 6, 1895–1903 (2017).

38. Englund, E., Schmidt, M., Nava, A. A., Lechner, A., Deng, K. et al. Expanding extender substrate selection for unnatural polyketide biosynthesis by acyltransferase domain exchange within a modular polyketide synthase. J. Am. Chem. Soc. 145, 8822–8832 (2023).

39. Kalkreuter, E., CroweTipton, J. M., Lowell, A. N., Sherman, D. H. & Williams, G. J. Engineering the substrate specificity of a modular polyketide synthase for installation of consecutive non-natural extender units. J. Am. Chem. Soc.141, 1961–1969 (2019).

40. Malico, A. A., Nichols, L. & Williams, G. J. Synthetic biology enabling access to designer polyketides. Curr. Opin. Chem. Biol. 58, 45–53 (2020).

41. Goranovič, D., Kosec, G., Mrak, P., Fujs, Š., Horvat, J., Kuščer, E., Kopitar, G. & Petković, H. Origin of the allyl group in FK506 biosynthesis. J. Biol. Chem. 285, 14292–14300 (2010).

42. Mo, S., Kim, D.-H., Lee, J. H., Oh, C.-H., Ban, Y.-H. et al. Biosynthesis of the allylmalonyl-CoA extender unit for the FK506 polyketide synthase proceeds through a dedicated polyketide synthase and facilitates the mutasynthesis of analogues. J. Am. Chem. Soc. 133, 976–985 (2011).

43. Jez, J. M., Bowman, M. E. & Noel, J. P. Expanding the biosynthetic repertoire of plant type III polyketide synthases by altering starter molecule specificity. Proc. Natl. Acad. Sci. U.S.A. 99, 5319–5324 (2002).

44. Yuzawa, S., Backman, T. W. H., Keasling, J. D. & Katz, L. Synthetic biology of polyketide synthases. J. Ind. Microbiol. Biotechnol. 45, 621–633 (2018).

45. Yuzawa, S., Deng, K., Wang, G., Baidoo, E. E. K., Northen, T. R. et al. Comprehensive in vitro analysis of acyltransferase domain exchanges in modular polyketide synthases and its application for short-chain ketone production. ACS Synth. Biol. 6, 139–147 (2017).

46. Buyachuihan, L., Reiners, S., Zhao, Y. & Grininger, M. The malonyl/acetyl-transferase from murine fatty acid synthase is a promiscuous engineering tool for editing polyketide scaffolds. Commun. Chem. 7, 187 (2024).

47. Kellenberger, L., Galloway, I. S., Sauter, G., Böhm, G., Hanefeld, U. et al. A polylinker approach to reductive loop swaps in modular polyketide synthases. ChemBioChem 9, 2740–2749 (2008).

48. Zargar, A., Lal, R., Valencia, L., Wang, J., Backman, T. W. H. et al. Chemoinformatic-guided engineering of polyketide synthases. J. Am. Chem. Soc. 142, 9896–9901 (2020).

49. Hagen, A., Poust, S., de Rond, T., Fortman, J. L., Katz, L. et al. Engineering a polyketide synthase for in vitro production of adipic acid. ACS Synth. Biol. 5, 21–27 (2016).

50. Montaño López, J., Duran, L. & Avalos, J. L. Physiological limitations and opportunities in microbial metabolic engineering. Nat Rev Microbiol 20, 35–48 (2022).

51. Rathnasingh, C., Raj, S. M., Lee, Y., Catherine, C., Ashok, S. et al. Production of 3-hydroxypropionic acid via malonyl-CoA pathway using recombinant *Escherichia coli* strains. J. Biotechnol. 157, 633–640 (2012).

52. Inui, M., Suda, M., Kimura, S., Yasuda, K., Suzuki, H. et al. Expression of *Clostridium acetobutylicum* butanol synthetic genes in *Escherichia coli*. Appl. Microbiol. Biotechnol. 77, 1305–1316 (2008).

53. Atsumi, S., Cann, A. F., Connor, M. R., Shen, C. R., Smith, K. M. et al. Metabolic engineering of *Escherichia coli* for 1-butanol production. Metab. Eng. 10, 305–311 (2008).

54. Atsumi, S., Hanai, T. & Liao, J. C. Non-fermentative pathways for synthesis of branched-chain higher alcohols as biofuels. Nature 451, 86–89 (2008).

55. Nielsen, D. R., Leonard, E., Yoon, S. H., Tseng, H. C., Yuan, C. et al. Engineering alternative butanol production platforms in heterologous bacteria. Metab. Eng. 11, 262–273 (2009).

56. Trinh, C. T., Li, J., Blanch, H. W. & Clark, D. S. Redesigning *Escherichia coli* metabolism for anaerobic production of isobutanol. Appl. Environ. Microbiol. 77, 4894–4904 (2011).

57. Kim, J. & Copley, S. D. Inhibitory cross-talk upon introduction of a new metabolic pathway into an existing metabolic network. Proc. Natl. Acad. Sci. U.S.A. 109, (2012).

58. Sirirungruang, S., Ad, O., Privalsky, T. M., Ramesh, S., Sax, J. L. et al. Engineering site-selective incorporation of fluorine into polyketides. Nat. Chem. Biol. 18, 886–893 (2022).

59. Rittner, A., Joppe, M., Schmidt, J. J., Mayer, L. M., Reiners, S. et al. Chemoenzymatic synthesis of fluorinated polyketides. Nat. Chem. 14, 1000–1006 (2022).

60. Buyachuihan, L., Reiners, S., Zhao, Y. & Grininger, M. The malonyl/acetyl-transferase from murine fatty acid synthase is a promiscuous engineering tool for editing polyketide scaffolds. Commun Chem 7, 187 (2024).

61. Müller, K., Faeh, C. & Diederich, F. Fluorine in pharmaceuticals: looking beyond intuition. Science 317, 1881–1886 (2007).

62. O’Hagan, D. & Deng, H. Enzymatic fluorination and biotechnological developments of the fluorinase. Chem. Rev. 115, 634–649 (2015).

63. Carbonell, P., Parutto, P., Baudier, C., Junot, C. & Faulon, J.-L. Retropath: automated pipeline for embedded metabolic circuits. ACS Synth. Biol. 3, 565–577 (2014).

64. Delépine, B., Duigou, T., Carbonell, P. & Faulon, J.-L. RetroPath2.0: a retrosynthesis workflow for metabolic engineers. Metab. Eng. 45, 158–170 (2018).

65. Hatzimanikatis, V., Li, C., Ionita, J. A., Henry, C. S., Jankowski, M. D. & Broadbelt, L. J. Exploring the diversity of complex metabolic networks. Bioinformatics 21, 1603–1609 (2005).

66. Gricourt, G., Meyer, P., Duigou, T. & Faulon, J.-L. Artificial intelligence methods and models for retro-biosynthesis: a scoping review. ACS Synth. Biol. 13, 2276–2294 (2024).

67. Yu, T., Boob, A. G., Volk, M. J., Liu, X., Cui, H. & Zhao, H. Machine learning-enabled retrobiosynthesis of molecules. Nat. Catal. 6, 137–151 (2023).

68. Koch, M., Duigou, T. & Faulon, J.-L. Reinforcement Learning for Bioretrosynthesis. ACS Synth. Biol. 9, 157–168 (2020).

69. Carbonell, P., Jervis, A. J., Robinson, C. J., Yan, C., Dunstan, M. et al. An automated design-build-test-learn pipeline for enhanced microbial production of fine chemicals. *Commun*. Biol. 1, 66 (2018).

70. Zheng, S., Zeng, T., Li, C., Chen, B., Coley, C. W., Yang, Y. & Wu, R. Deep learning driven biosynthetic pathways navigation for natural products with BioNavi-NP. Nat. Commun. 13, 3342 (2022).

71. Levin, I., Liu, M., Voigt, C., Coley, C. Merging enzymatic and synthetic chemistry with computational synthesis planning. Nat. Commun. 13, 7747 (2022).

72. Finnigan, W., Hepworth, L. J., Flitsch, S. L. & Turner, N. J. RetroBioCat as a computer-aided synthesis planning tool for biocatalytic reactions and cascades. Nat Catal 4, 98–104 (2021).

73. Chen, S. & Jung, Y. Deep retrosynthetic reaction prediction using local reactivity and global attention. JACS Au 1, 1612–1620 (2021).

74. Dong, J., Zhao, M., Liu, Y., Su, Y. & Zeng, X. Deep learning in retrosynthesis planning: datasets, models and tools. Briefings in Bioinformatics 23, bbab391 (2022).

75. Upadhyay, V., Anand, M. & Maranas, C. D. novoStoic2.0: An integrated framework for pathway synthesis, thermodynamic evaluation, and enzyme selection. Preprint at 10.1101/2024.09.27.615368 (2024).

76. Dejong, C. A., Chen, G. M., Li, H., Johnston, C. W., Edwards, M. R., Rees, P. N., Skinnider, M. A. et al. Polyketide and nonribosomal peptide retro-biosynthesis and global gene cluster matching. Nat. Chem. Biol. 12, 1007–1014 (2016).

77. Skinnider, M. A., Johnston, C. W., Gunabalasingam, M., Merwin, N. J., Kieliszek, A. M. et al. Comprehensive prediction of secondary metabolite structure and biological activity from microbial genome sequences. Nat. Commun. 11, 6058 (2020).

78. Blin, K., Shaw, S., Kloosterman, A. M., Charlop-Powers, Z., van Wezel, G. P., Medema, M. H. & Weber, T. antiSMASH 6.0: improving cluster detection and comparison capabilities. Nucleic Acids Res. 49, W29–W35 (2021).

79. Terlouw, B. R., Blin, K., Navarro-Muñoz, J. C., Avalon, N. E., Chevrette, M. G. et al. MIBiG 3.0: a community-driven effort to annotate experimentally validated biosynthetic gene clusters. Nucleic Acids Res. 51, D603–D610 (2023).

80. Kautsar, S. A., Suarez Duran, H. G., Blin, K., Osbourn, A. & Medema, M. H. plantiSMASH: automated identification, annotation and expression analysis of plant biosynthetic gene clusters. Nucleic Acids Res. 45, W55–W63 (2017).

81. Beber, M. E., Gollub, M. G., Mozaffari, D., Shebek, K. M., Flamholz, A. I. et al. eQuilibrator 3.0: a database solution for thermodynamic constant estimation. Nucleic Acids Res. 50, D603–D609 (2022).

82. Wang, L., Upadhyay, V. & Maranas, C. D. dGPredictor: Automated fragmentation method for metabolic reaction free energy prediction and de novo pathway design. PLoS Comput Biol 17, e1009448 (2021).

83. Zhang, Y., Yang, Y., & Zhang, Y. α-Pyrones and their derivatives from two *Cryptocarya* species. Phytochemistry 39, 481–485 (1995).

84. Iinuma, H., Nakamura, H., Naganawi, H., Masuda, T., Takano, S. et al. Basidalin, a new antibiotic from basidiomycetes. J. Antibiot. 36, 448–450 (1983).

85. Tao, X.B., LaFrance, S., Xing, Y., Nava, A.A., Garcia Martin, H. et al. ClusterCAD 2.0: an updated computational platform for chimeric type I polyketide synthase and nonribosomal peptide synthetase design. Nucleic Acids Res. 51, D532–D537 (2023).

86. Ni, Z., Stine, A. E., Tyo, K. E. J. & Broadbelt, L. J. Curating a comprehensive set of enzymatic reaction rules for efficient novel biosynthetic pathway design. Metab. Eng. 65, 79–87 (2021).

87. Ni, Z. Data-Driven Approaches to Predict and Utilize Enzyme Promiscuity for Novel Biosynthesis Design. ProQuest Dissertations and Theses (Northwestern University, United States -- Illinois, 2022).

88. Caspi, R., Billington, R., Keseler, I.M., Kothari, A., Krummenacker, M. et al. The MetaCyc database of metabolic pathways and enzymes – a 2019 update. Nucleic Acids Res. 48, D445–D453 (2020).

89. Kanehisa, M., Furumichi, M., Tanabe, M., Sato, Y. & Morishima, K. KEGG: new perspectives on genomes, pathways, diseases and drugs. Nucleic Acids Res. 45, D353–D361 (2017).

90. Schomburg, I., Jeske, L., Ulbrich, M., Placzek, S., Chang, A. et al. The BRENDA enzyme information system–from a database to an expert system. J. Biotechnol. 261, 194–206 (2017).

91. Chainani, Y., Ni, Z., Shebek, K. M., Broadbelt, L. J. & Tyo, K. E. J. DORA-XGB: an improved enzymatic reaction feasibility classifier trained using a novel synthetic data approach. Mol. Syst. Des. Eng. 10, 129–142 (2025).

92. Robbins, T., Kapilivsky, J., Cane, D. E. & Khosla, C. Roles of conserved active site residues in the ketosynthase domain of an assembly line polyketide synthase. Biochemistry 55, 4476–4484 (2016).

93. Dan, Q., Chiu, Y., Lee, N., Pereira, J. H., Rad, B. et al. A polyketide-based biosynthetic platform for diols, amino alcohols and hydroxy acids. Nat. Catal. 2025, 1–15 (2025).

94. Choi, S., Kim, H. U., Kim, T. Y., Kim, W. J., Lee, M. H. et al. Production of 4-hydroxybutyric acid by metabolically engineered *Mannheimia succiniciproducens* and its conversion to γ-butyrolactone by acid treatment. Metab. Eng. 20, 73–83 (2013).

95. Choi, S., Kim, H. U., Kim, T. Y. & Lee, S. Y. Systematic engineering of TCA cycle for optimal production of a four-carbon platform chemical 4-hydroxybutyric acid in Escherichia coli. Metabolic Engineering 38, 264–273 (2016).

96. Hein, S., Söhling, B., Gottschalk, G. & Steinbüchel, A. Biosynthesis of poly(4-hydroxybutyric acid) by recombinant strains of Escherichia coli1. FEMS Microbiology Letters 153, 411–418 (2006).

97. Min, K., Kim, S., Yum, T., Kim, Y., Sang, B.-I. et al. Conversion of levulinic acid to 2-butanone by acetoacetate decarboxylase from *Clostridium acetobutylicum*. Appl. Microbiol. Biotechnol. 97, 5627–5634 (2013).

98. Fadda, S., Lebert, A., Leroy-Sétrin, S. & Talon, R. Decarboxylase activity involved in methyl ketone production by *Staphylococcus carnosus* 833, a strain used in sausage fermentation. FEMS Microbiol. Lett. 210, 209–214 (2002).

99. Park, J., Rodríguez-Moyá, M., Li, M., Pichersky, E., San, K. Y. et al. Synthesis of methyl ketones by metabolically engineered *Escherichia coli*. J. Ind. Microbiol. Biotechnol. 39, 1703–1712 (2012).

100. Lee, H., DeLoache, W. C. & Dueber, J. E. Spatial organization of enzymes for metabolic engineering. Metab. Eng. 14, 242–251 (2012).

101. Sun, X., Yuan, Y., Chen, Q., Nie, S., Guo, J. et al. Metabolic pathway assembly using docking domains from type I cis-AT polyketide synthases. Nat. Commun. 13, 5541 (2022).

102. Vaithegi, K. & Prasad, K. R. Total synthesis of (−)-cryptofolione. Tetrahedron 79, 131842 (2021)..

103. Trefzer, A., Pelzer, S., Schimana, J., Stockert, S., Bihlmaier, C. et al. Biosynthetic gene cluster of simocyclinone, a natural multihybrid antibiotic. Antimicrob. Agents Chemother. 46, 1174–1182 (2002).

104. Kakavas, S. J., Katz, L. & Stassi, D. Identification and characterization of the niddamycin polyketide synthase genes from *Streptomyces caelestis*. J. Bacteriol. 179, 7515–7522 (1997).

105. Acosta, J. A., Muddala, R., Barbosa, L. C. & Boukouvalas, J. Total synthesis of the antitumor antibiotic basidalin. J. Org. Chem. 81, 6883–6886 (2016).

106. Keatinge-Clay, A. T. Stereocontrol within polyketide assembly lines. Nat. Prod. Rep. 33, 141–149 (2016).

107. Weissman, K. J. Polyketide stereocontrol: a study in chemical biology. Beilstein J. Org. Chem. 13, 348–371 (2017).

108. Hall, M. Enzymatic strategies for asymmetric synthesis. RSC Chemical Biology 2, 958–989 (2021).

109. Heid, E., Probst, D., Green, W. H. & Madsen, G. K. EnzymeMap: curation, validation and data-driven prediction of enzymatic reactions. Chem. Sci. 14, 14229–14242 (2023).

110. Duigou, T., du Lac, M., Carbonell, P. & Faulon, J.-L. RetroRules: a database of reaction rules for engineering biology. Nucleic Acids Res. 47, D1229–D1235 (2019).

111. Wiest, O., Bauer, C., Helquist, P., Norrby, P. O. & Genheden, S. Finding relevant retrosynthetic disconnections for stereocontrolled reactions. J. Chem. Inf. Model. 64, 5796–5805 (2024).

112. Upadhyay, V., Boorla, V. S. & Maranas, C. D. Rank-ordering of known enzymes as starting points for re-engineering novel substrate activity using a convolutional neural network. Metab. Eng. 78, 171–182 (2023).

113. Carbonell, P., Wong, J., Swainston, N., Takano, E., Turner, N. J. et al. Selenzyme: enzyme selection tool for pathway design. Bioinformatics 34, 2153–2154 (2018).

114. Dalkiran, A., Rifaioglu, A. S., Martin, M. J., Cetin-Atalay, R., Atalay, V. et al. ECPred: a tool for the prediction of the enzymatic functions of protein sequences based on the EC nomenclature. BMC Bioinformatics 19, 1–13 (2018).

115. Yu, T., Cui, H., Li, J. C., Luo, Y., Jiang, G. et al. Enzyme function prediction using contrastive learning. Science 379, 1358–1363 (2023).

116. Chowdhury, A. & Maranas, C. D. Designing overall stoichiometric conversions and intervening metabolic reactions. Sci. Rep. 5, 16009 (2015).

117. Porokhin, V., Amin, S. A., Nicks, T. B., Gopinarayanan, V. E., Nair, N. U. et al. Analysis of metabolic network disruption in engineered microbial hosts due to enzyme promiscuity. Metab. Eng. Commun. 12, e00170 (2021).

118. Arnold, F. H. Design by directed evolution. Acc. Chem. Res. 31, 125–131 (1998).

119. Wang, Y., Xue, P., Cao, M., Yu, T., Lane, S. T., et al. Directed evolution: methodologies and applications. Chem. Rev. 121, 12384–12444 (2021).

120. Dauparas, J., Anishchenko, I., Bennett, N., Bai, H., Ragotte, R. J. et al. Robust deep learning–based protein sequence design using ProteinMPNN. Science 378, 49–56 (2022).

121. Watson, J. L., Juergens, D., Bennett, N. R., Trippe, B. L., Yim, J. et al. De novo design of protein structure and function with RFdiffusion. Nature 620, 1089–1100 (2023).

122. Nava, A. A., Roberts, J., Haushalter, R. W., Wang, Z. & Keasling, J. D. Module-based polyketide synthase engineering for *de novo* polyketide biosynthesis. ACS Synth. Biol. 12, 3148–3155 (2023).

123. Jumper, J., Evans, R., Pritzel, A., Green, T., Figurnov, M. et al. Highly accurate protein structure prediction with AlphaFold. Nature 596, 583–589 (2021).

124. Orsi, M. & Reymond, J.-L. One chiral fingerprint to find them all. J. Cheminform. 16, 53 (2024).

125. Gobbi, A., Giannetti, A. M., Chen, H. & Lee, M. L. Atom-atom-path similarity and sphere exclusion clustering: tools for prioritizing fragment hits. J. Cheminform. 7, 1–11 (2015).

126. Carhart, R. E., Smith, D. H. & Venkataraghavan, R. Atom pairs as molecular features in structure–activity studies: definition and applications. J. Chem. Inf. Comput. Sci. 25, 64–73 (1985).

127. Mabesoone, M. F., Leopold-Messer, S., Minas, H. A., Chepkirui, C., Chawengrum, P. et al. Evolution-guided engineering of trans-acyltransferase polyketide synthases. Science 383, 1312–1317 (2024).

128. Shebek, K. M., Strutz, J., Broadbelt, L. J. & Tyo, K. E. Pickaxe: a Python library for the prediction of novel metabolic reactions. BMC Bioinformatics 24, 106 (2023).

129. The UniProt Consortium et al. UniProt: the Universal Protein Knowledgebase in 2025. Nucleic Acids Research 53, D609–D617 (2025).

130. Noor, E., Bar-Even, A., Flamholz, A., Reznik, E., Liebermeister, W. et al. Pathway thermodynamics highlights kinetic obstacles in central metabolism. PLoS Comput. Biol. 10, e1003483 (2014).

