## Supporting Information for "*BioPKS Pipeline*: An integrated platform for merging the computational design of chimeric type I polyketide synthases with enzymatic pathways for chemical biosynthesis"

### Supplementary Information

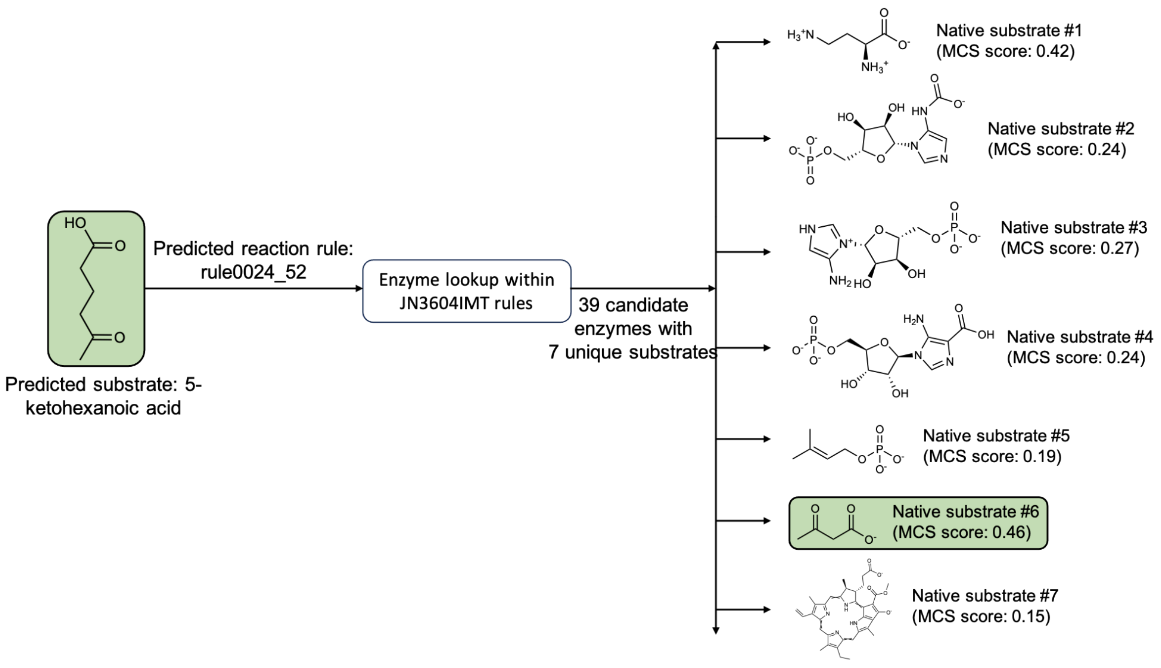


**SI Fig. 1. Selection of enzymes for predicted decarboxylation of 5-ketohexanoic acid to produce 2-pentanone.**

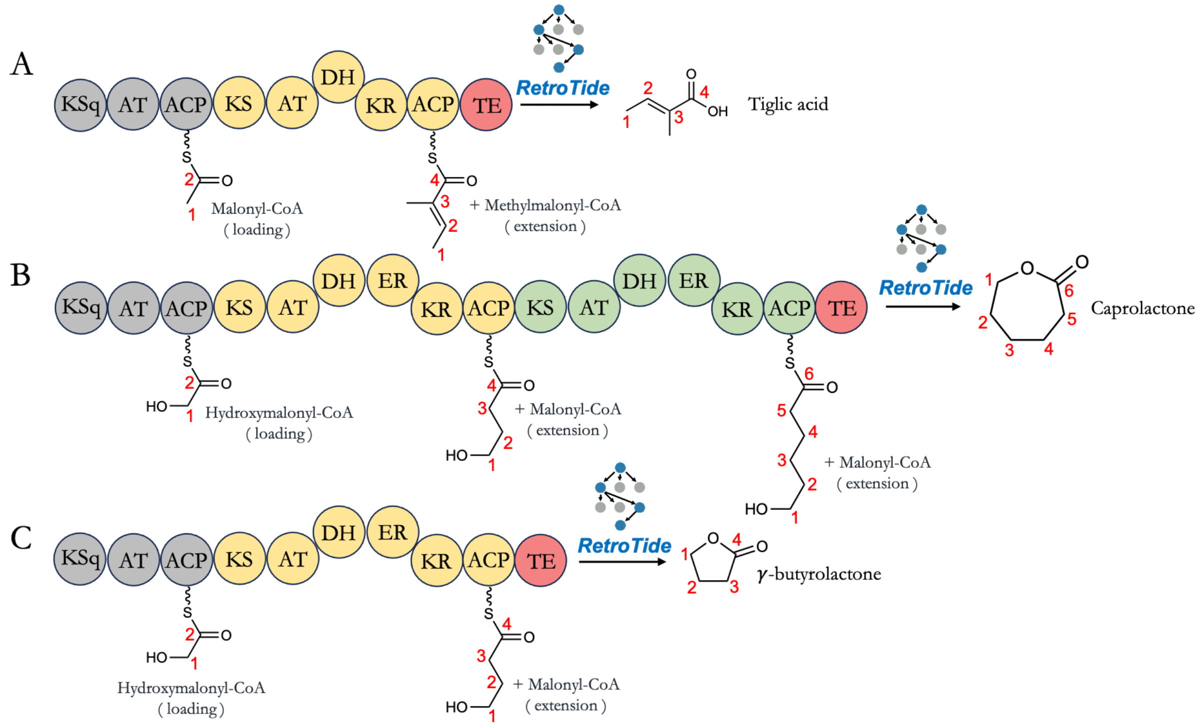


**SI Fig. 2. Proposed biosynthesis of (A) tiglic acid, (B) caprolactone, and (C) γ–butyrolactone by BioPKS Pipeline.**

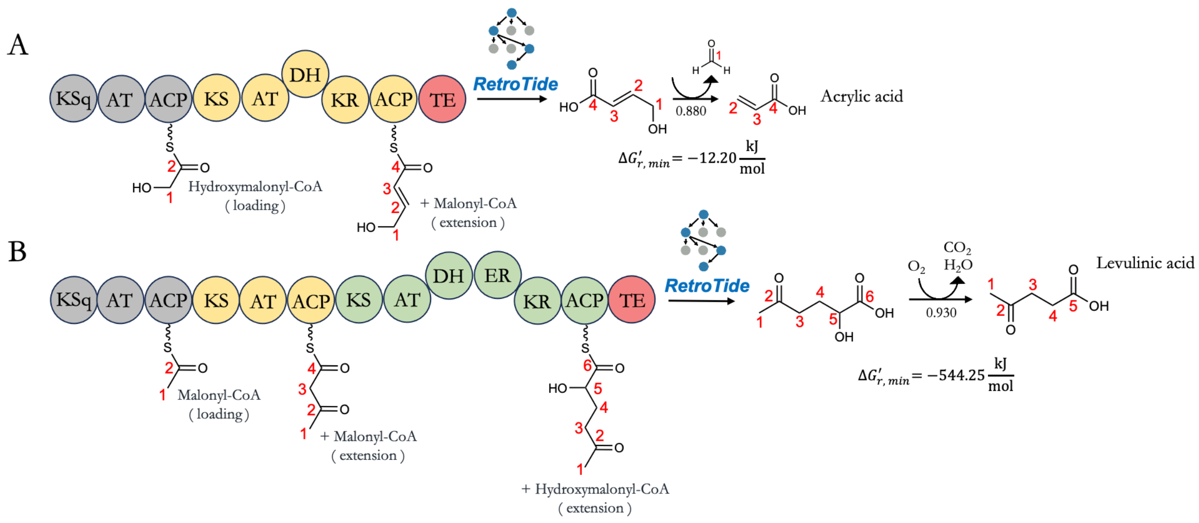


**SI Fig. 3. Successfully proposed biosynthesis of (A) acrylic acid, (B) levulinic acid by BioPKS Pipeline.**

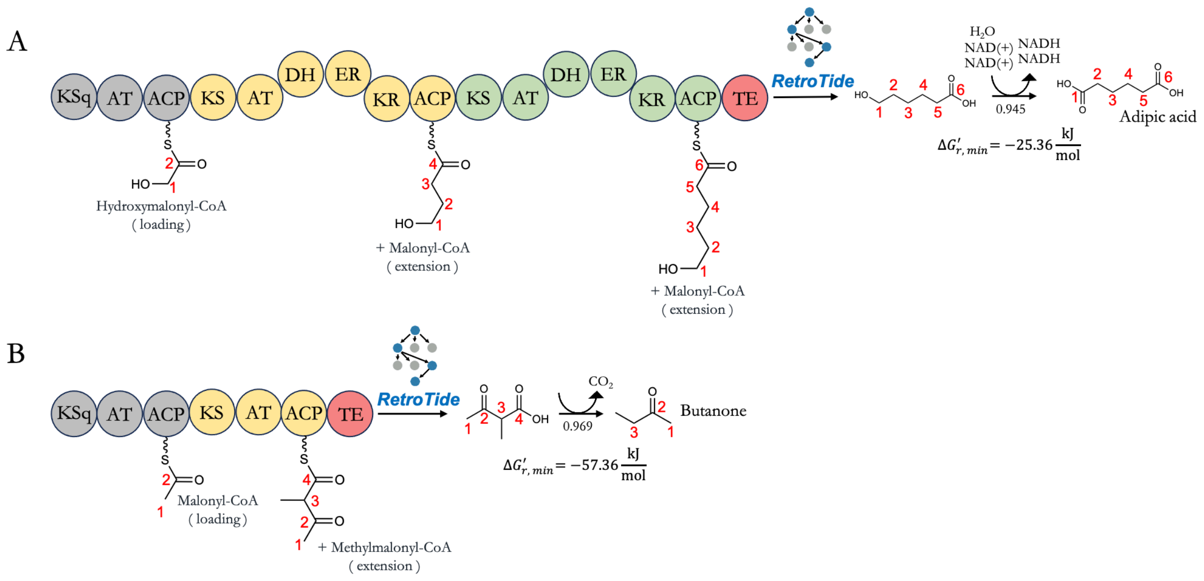


**SI Fig. 4. Successfully proposed biosynthesis of (A) adipic acid and (B) butanone by BioPKS Pipeline.**

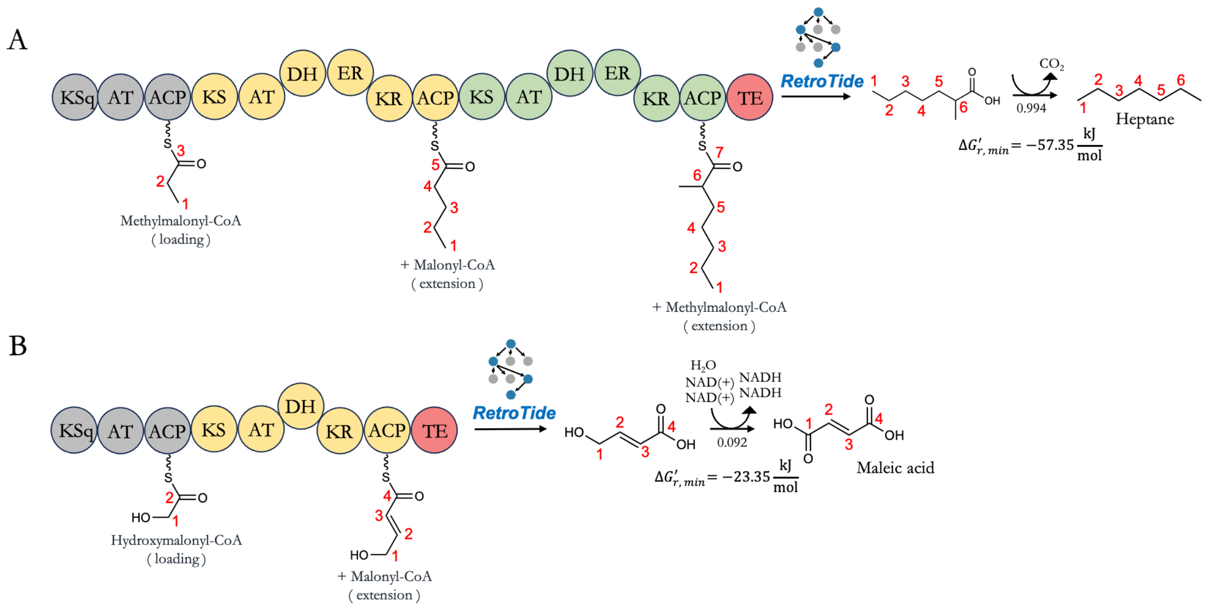


**SI Fig. 5. Successfully proposed biosynthesis of (A) heptane and (B) maleic acid by BioPKS Pipeline.**

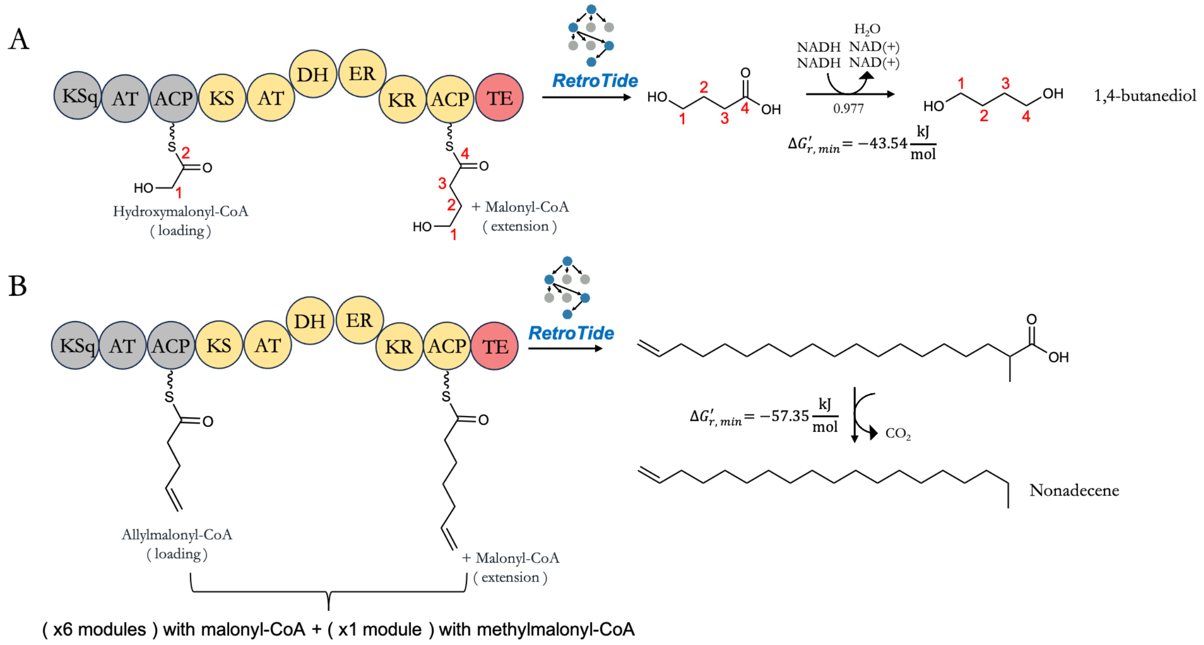


**SI Fig. 6. Successfully proposed biosynthesis of (A) 1,4-butanediol and (B) nonadecene by BioPKS Pipeline.**

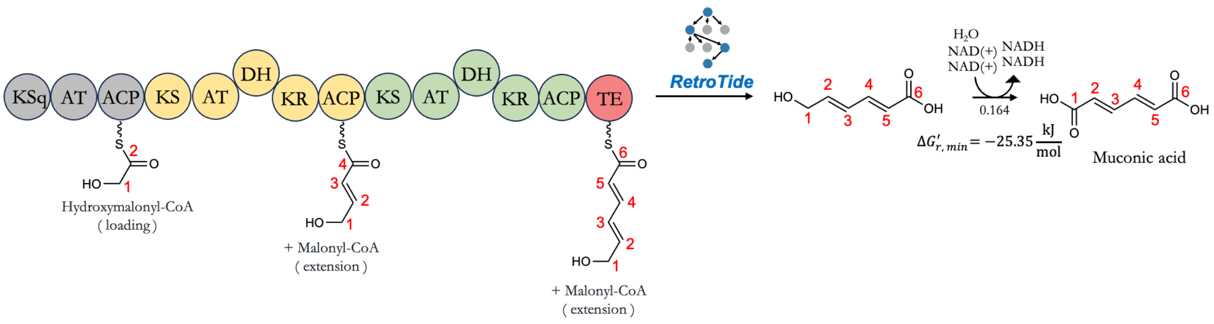


**SI Fig. 7. Successfully proposed biosynthesis of (A) muconic acid by BioPKS Pipeline.**

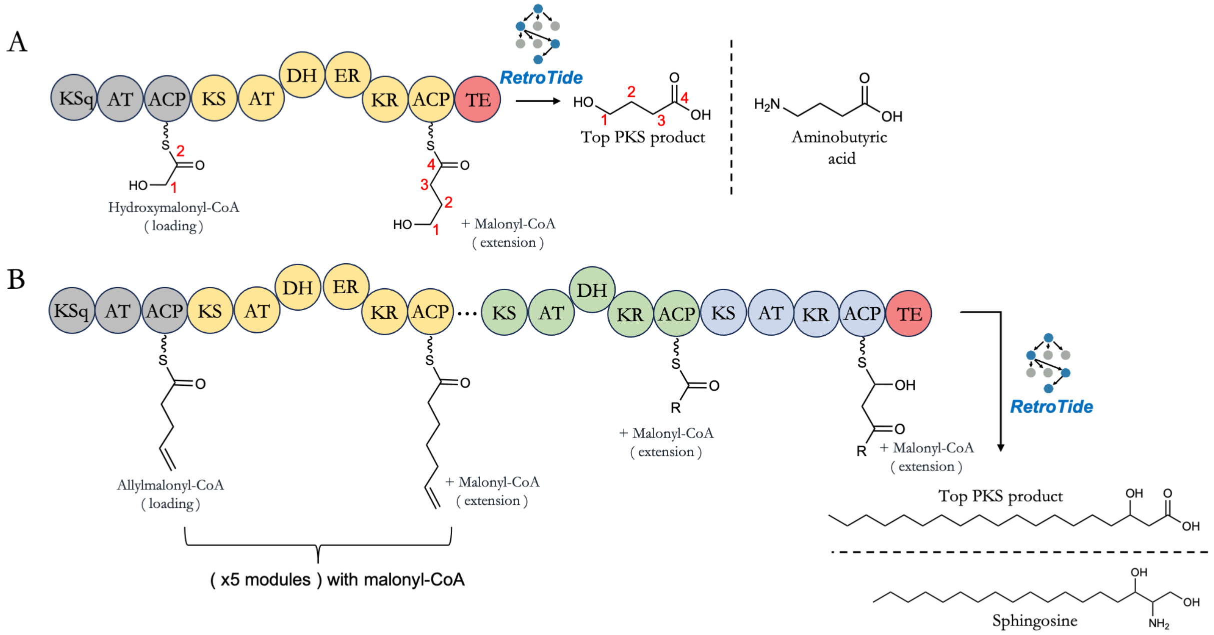


**SI Fig. 8. Unsuccessfully proposed biosynthesis of (A) aminobutyric-acid and (B) sphingosine by BioPKS Pipeline (top-ranked PKS designs shown).**

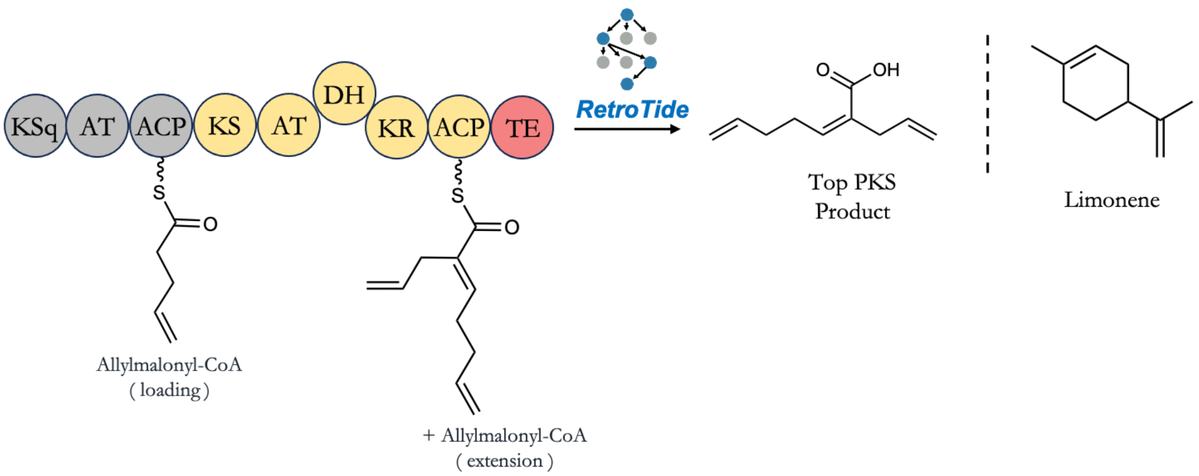


**SI Fig. 9. Unsuccessfully proposed biosynthesis of limonene by BioPKS Pipeline (top-ranked PKS design shown). For limonene specifically, even after attempting more than two post-PKS modifications, the target structure of limonene was not reached.**

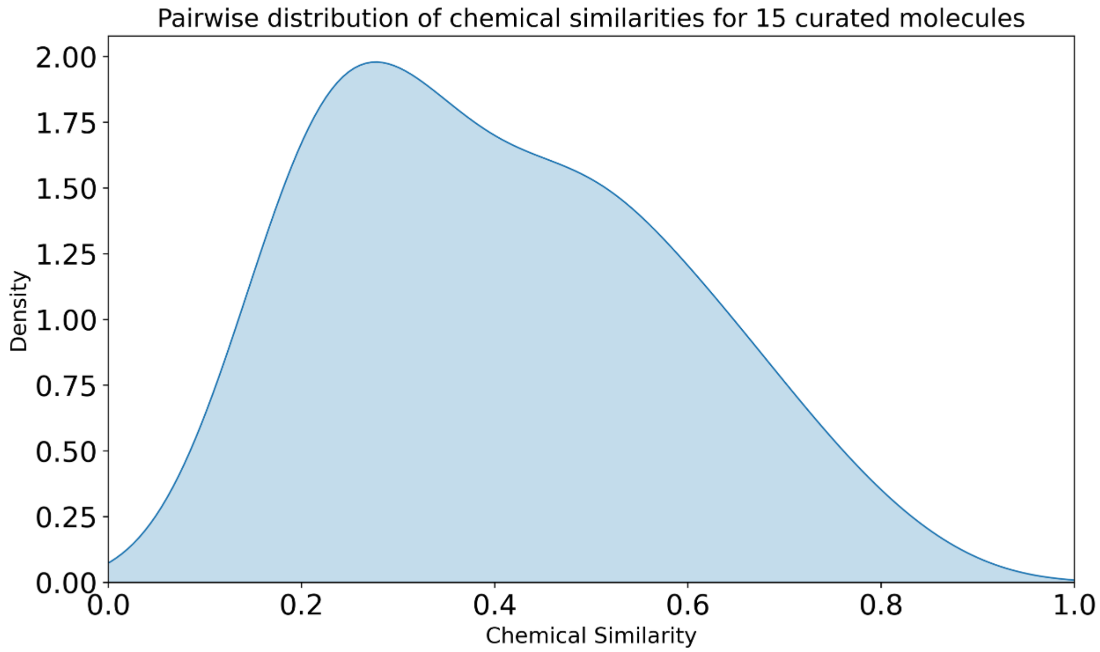


**SI Fig. 10. Pairwise chemical similarity distribution of 15 curated molecules for the initial prototyping of BioPKS Pipeline.**

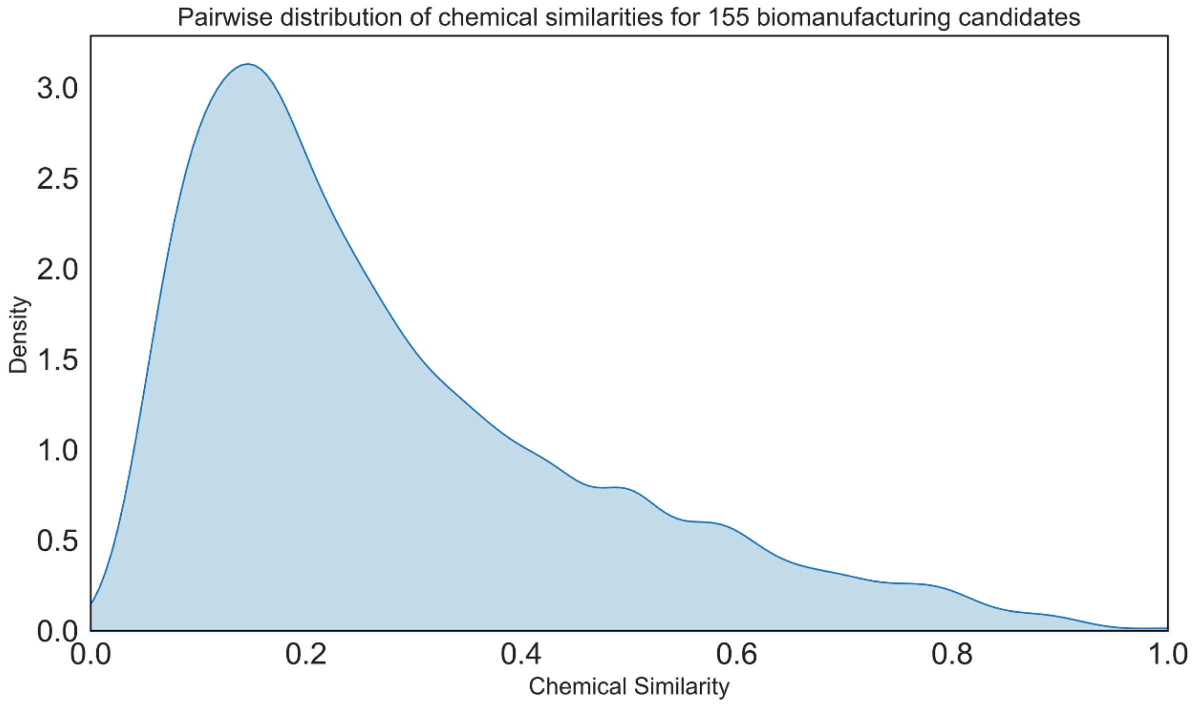


**SI Fig. 11. Pairwise chemical similarity distribution of 155 biomanufacturing candidates for the validation of BioPKS Pipeline.**

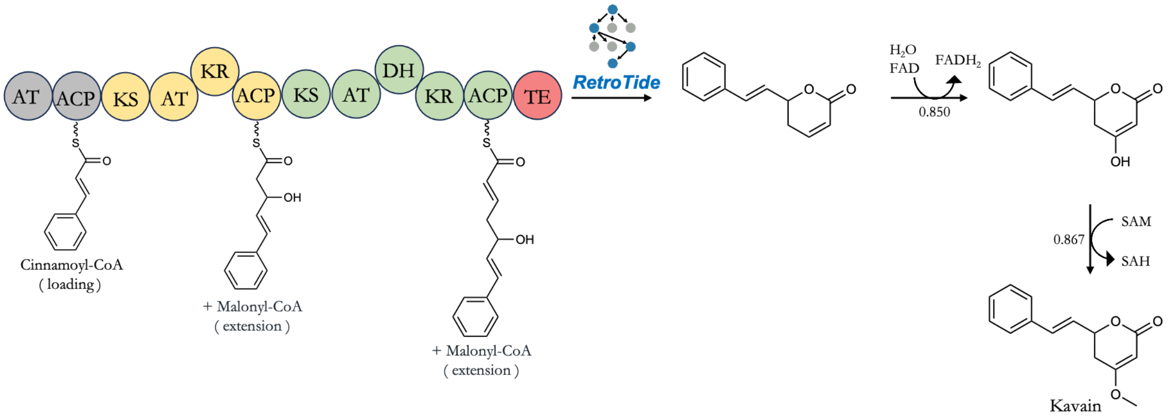


**SI Fig. 12. Successfully proposed biosynthesis of kavain by BioPKS Pipeline.**

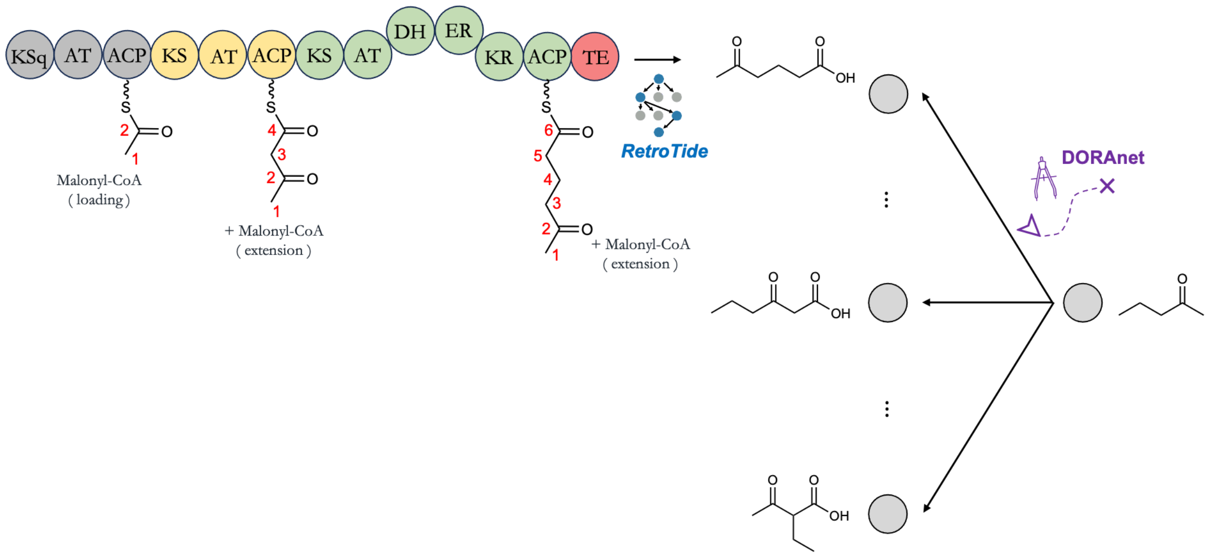


**SI Fig. 13. Reverse synthesis of 2-Pentanone by running DORAnet backwards and RetroTide forward.**

**SI Table 1. List of PKS starter units currently available for use in RetroTide**

| # | Name | SMILES |
|---|---|---|
| 1 | Acetyl- | CC(=O)[S] |
| 2 | Propionyl- | CCC(=O)[S] |
| 3 | Malonyl- | CC(=O)[S] |
| 4 | Methylmalonyl- | CCC(=O)[S] |
| 5 | Allylmalonyl- | C=CCCC([S])=O |
| 6 | Methoxymalonyl- | COCC(=O)[S] |
| 7 | Hydroxymalonyl- | [S]C(CO)=O |
| 8 | Succinyl- | [S]C(CCC(O)=O)=O |
| 9 | Butyrylmalonyl- | CCCCCC([S])=O |

| | | |
|---|---|---|
| 10 | Isobutyrylmalonyl- | CC(C)C(=O)[S] |
| 11 | 2-methylbutyrylmalonyl- | CCC(C)C(=O)[S] |
| 12 | DCP | [S]C(C1=CNC(Cl)=C1Cl)=O |
| 13 | Cemal- | CC(=O)[S] |
| 14 | CHC- | C1CCCCC1C(=O)[S] |
| 15 | trans-1,2-CPDA | C1CC[C@@H](C(=O)O)[C@@H]1C(=O)[S] |
| 16 | Cyclopentene | C1(=O)C(=CCC1)C(=O)[S] |
| 17 | Pyr- | P[S]C(C1=CC=CN1)=O |
| 18 | Cinnamoyl- | O=C([S])/C=C/C1=CC=CC=C1 |
| 19 | ABHA- | [S]C(C1=CC(O)=CC(N)=C1)=O |
| 20 | Isovalerate- | CC(CC([S])=O)C |
| 21 | PABA- | NC1=CC=C(C([S])=O)C=C1 |
| 22 | Guan | NC(NCC([S])=O)=[NH2+] |
| 23 | Mthz | CC1=NC(C([S])=O)=CS1 |
| 24 | DHCH | O[C@H]1[C@H](O)CCC(C([S])=O)C1 |
| 25 | DHCHene | O[C@H]1[C@H](O)CC=C(C([S])=O)C1 |
| 26 | plac- | O=C([S])CC1=CC=CC=C1 |
| 27 | ben | [S]C(C1=CC=CC=C1)=O |

| | | |
|---|---|---|
| 28 | PNBA | [S]C(C1=CC=C([N+]([O-])=O)C=C1)=O |
| 29 | ema | [S]C([C@@H](CC)C(N)=O)=O |
| 30 | 3measp | [S]C([C@@H](C)CNC([C@@H](N)C)=O)=O |

**SI Table 2. List of PKS extender units currently available for use in RetroTide**

| # | Name | SMILES |
|---|---|---|
| 1 | Malonyl- | O=C(O)CC(=O)[S] |
| 2 | Methylmalonyl- | C[C@@H](C(=O)O)C(=O)[S] |
| 3 | Allylmalonyl- | C=CC[C@@H](C(=O)O)C(=O)[S] |
| 4 | Methoxymalonyl- | CO[C@@H](C(=O)O)C(=O)[S] |
| 5 | Ethylmalonyl- | CC[C@@H](C(=O)O)C(=O)[S] |
| 6 | Butyrylmalonyl- | CCCC[C@@H](C(=O)O)C(=O)[S] |
| 7 | Hydroxymalonyl- | OC(C(C([S])=O)O)=O |
| 8 | Isobutyrylmalonyl- | [S]C([C@@H](C(O)=O)CC(C)C)=O |
| 9 | D-isobutyrylmalonyl- | [S]C([C@H](C(O)=O)CC(C)C)=O |
| 10 | DCP- | ClC1=C(Cl)NC=C1CCCCC(C(O)=O)C([S])=O |
| 11 | Hexylmalonyl- | CCCCCC[C@@H](C(=O)O)C(=O)[S] |

**SI Table 3. Biomanufacturing candidates used to report BioPKS Pipeline's performance.**

| # | Name | SMILES | Synthesis |
|---|---|---|---|
| 1 | Propionaldehyde | CCC=O | Reached in both 1 post-PKS and 2 post-PKS steps |
| 2 | Cyclohexanol | OC1CCCCC1 | Not reached |
| 3 | Ethyne | C#C | Not reached |
| 4 | Ascorbic acid | O=C1OC(C(O)CO)C(O)=C1O | Not reached |
| 5 | Citric acid | O=C(O)CC(O)(CC(=O)O)C(=O)O | Not reached |
| 6 | 2-Methyl-1-propanol | CC(C)CO | Reached in 2 post-PKS steps only |
| 7 | Glutaric acid | C(CC(=O)O)CC(=O)O | Reached in 2 post-PKS steps only |
| 8 | Camphene | CC1(C2CCC(C2)C1=C)C | Not reached |
| 9 | Maleate | O=C(O)C=CC(O)=O | Reached in both 1 post-PKS and 2 post-PKS steps |
| 10 | Acrylic acid | C=CC(=O)O | Reached in both 1 post-PKS and 2 post-PKS steps |
| 11 | Methyl acetate | CC(=O)OC | Reached in 2 post-PKS steps only |
| 12 | Ethanal | CC=O | Reached in both 1 post-PKS and 2 post-PKS steps |
| 13 | 6-Hexanolide | O=C1CCCCCO1 | Reached simply with PKSs |
| 14 | 2-Methylbutyraldehyde | CCC(C)C=O | Reached in both 1 post-PKS and 2 post-PKS steps |
| 15 | Propene | CC=C | Reached in both 1 post-PKS and 2 post-PKS steps |
| 16 | D-gluconate | O=C(O)C(O)C(O)C(O)C(O)CO | Reached simply with PKSs |
| 17 | Propenenitrile | C=CC#N | Reached in 2 post-PKS steps only |
| 18 | Z-3,7-Dimethylocta-2,6-dien-1-ol | CC(C)=CCCC(C)=CC(O)=O | Not reached |
| 19 | Caprylic acid | CCCCCCCC(=O)O | Reached in both 1 post-PKS and 2 post-PKS steps |
| 20 | 2-Propanol | CC(C)O | Reached in both 1 post-PKS and 2 post-PKS steps |
| 21 | 1,2-Ethanediol | OCCO | Reached in both 1 post-PKS and 2 post-PKS steps |
| 22 | Butanoic acid | CCCC(=O)O | Reached in both 1 post-PKS and 2 post-PKS steps |
| 23 | 2_Butenoic_acid | CC=CC(=O)O | Reached in both 1 post-PKS and 2 post-PKS steps |
| 24 | Acetic acid | CC(=O)O | Reached in both 1 post-PKS and 2 post-PKS steps |
| 25 | Propanoic acid | CCC(=O)O | Reached in both 1 post-PKS and 2 post-PKS steps |
| 26 | NN_dimethylmethanamide | CN(C)C=O | Not reached |

| | | | |
|---|---|---|---|
| 27 | Glyoxylate | C(=O)C(=O)O | Reached in both 1 post-PKS and 2 post-PKS steps |
| 28 | 1-Amino-2-propanol | CC(CN)O | Reached in 2 post-PKS steps only |
| 29 | Stearate | CCCCCCCCCCCCCCCCCC(O)=O | Reached in 2 post-PKS steps only |
| 30 | Hexadecanoate | CCCCCCCCCCCCCCCC(O)=O | Reached in 2 post-PKS steps only |
| 31 | 2-Propenamide | C=CC(=O)N | Reached in 2 post-PKS steps only |
| 32 | beta-Mycrene | CC(=CCCC(=C)C=C)C | Not reached |
| 33 | Cyclohexanamine | C1CCC(CC1)N | Not reached |
| 34 | Adipic acid | C(CCC(=O)O)CC(=O)O | Reached in both 1 post-PKS and 2 post-PKS steps |
| 35 | L-menthol | CC1CCC(C(C1)O)C(C)C | Not reached |
| 36 | Limonene | C=C(C)C1CC=C(C)CC1 | Not reached |
| 37 | Gluconic lactone | C(C1C(C(C(C(=O)O1)O)O)O)O | Reached in both 1 post-PKS and 2 post-PKS steps |
| 38 | Linalool | CC(C)=CCCC(C=C)(O)C | Not reached |
| 39 | Acetylacetone | CC(=O)CC(C)=O | Reached in both 1 post-PKS and 2 post-PKS steps |
| 40 | Succinic_acid | C(CC(=O)O)C(=O)O | Reached in both 1 post-PKS and 2 post-PKS steps |
| 41 | Dodecanoate | CCCCCCCCCCCC(O)=O | Reached in both 1 post-PKS and 2 post-PKS steps |
| 42 | Oxalic_acid | C(=O)(C(=O)O)O | Reached in both 1 post-PKS and 2 post-PKS steps |
| 43 | 2_Dehydro_D_gluconate | O=C(C(O)=O)C(O)C(O)C(O)CO | Reached in 2 post-PKS steps only |
| 44 | Propanol | CCCO | Reached in both 1 post-PKS and 2 post-PKS steps |
| 45 | Prenyl alcohol | CC(=CCO)C | Reached in 2 post-PKS steps only |
| 46 | Methyl_methacrylate | CC(=C)C(=O)OC | Not reached |
| 47 | CH3N | CN(C)C | Not reached |
| 48 | Ethyl methyl ketone | CCC(=O)C | Reached in both 1 post-PKS and 2 post-PKS steps |
| 49 | Z_Octadec_9_enoic_acid | CCCCCCCCC=CCCCCCCCC(=O)O | Not reached |
| 50 | Itaconic acid | C=C(CC(=O)O)C(=O)O | Reached in 2 post-PKS steps only |
| 51 | CH32NH | CNC | Not reached |
| 52 | Oxaldehyde | C(=O)C=O | Reached in 2 post-PKS steps only |
| 53 | Fumaric acid | C(=CC(=O)O)C(=O)O | Reached in both 1 post-PKS and 2 post-PKS steps |
| 54 | alpha_pinene | CC1=CCC2CC1C2(C)C | Not reached |
| 55 | Butyraldehyde | CCCC=O | Reached in both 1 post-PKS and 2 post-PKS steps |
| 56 | Isoprene | CC(=C)C=C | Reached in 2 post-PKS steps only |
| 57 | Trolamine | C(CO)N(CCO)CCO | Not reached |

| | | | |
|---|---|---|---|
| 58 | Glycolic acid | C(C(=O)O)O | Reached in both 1 post-PKS and 2 post-PKS steps |
| 59 | 3_Methylbut_2_enal | CC(=CC=O)C | Reached in 2 post-PKS steps only |
| 60 | Aminoethanol | NCCO | Reached in 2 post-PKS steps only |
| 61 | Hexadecanol | CCCCCCCCCCCCCCCCO | Not reached |
| 62 | Cane sugar | C(C1C(C(C(C(O1)OC2(C(C(C(O2)CO)O)O)CO)O)O)O)O | Not reached |
| 63 | Acetone | CC(=O)C | Reached in both 1 post-PKS and 2 post-PKS steps |
| 64 | Ethanol | CCO | Reached in both 1 post-PKS and 2 post-PKS steps |
| 65 | 12_Propanediol | CC(CO)O | Reached in both 1 post-PKS and 2 post-PKS steps |
| 66 | Alphahydroxy-isobutyronitrile | CC(C)(C#N)O | Reached in 2 post-PKS steps only |
| 67 | Oleyl_amide | CCCCCCCCC=CCCCCCCCC(=O)N | Not reached |
| 68 | D-Mannitol | OCC(O)C(O)C(O)C(O)CO | Reached in both 1 post-PKS and 2 post-PKS steps |
| 69 | Linoleic acid | CCCCCC=CCC=CCCCCCCCC(O)=O | Not reached |
| 70 | Decanoate | CCCCCCCCCC(O)=O | Reached in both 1 post-PKS and 2 post-PKS steps |
| 71 | Tetradecanoate | CCCCCCCCCCCCCC(O)=O | Reached in 2 post-PKS steps only |
| 72 | Glycerin | C(C(CO)O)O | Reached in both 1 post-PKS and 2 post-PKS steps |
| 73 | Cyclohexanone | C1CCC(=O)CC1 | Not reached |
| 74 | Cyclohexane | C1CCCCC1 | Not reached |
| 75 | Butanol | CCCCO | Reached in both 1 post-PKS and 2 post-PKS steps |
| 76 | Hexanol | CCCCCCO | Reached in both 1 post-PKS and 2 post-PKS steps |
| 77 | Malic_acid | C(C(C(=O)O)O)C(=O)O | Reached in both 1 post-PKS and 2 post-PKS steps |
| 78 | Ethanamine | CCN | Reached in 2 post-PKS steps only |
| 79 | D_Fructose | C1C(C(C(C(O1)(CO)O)O)O)O | Reached in 2 post-PKS steps only |
| 80 | Cyclopentanone | C1CCC(=O)C1 | Not reached |
| 81 | Isobutyric acid | CC(C)C(=O)O | Reached in 2 post-PKS steps only |
| 82 | Caprolactam | C1CCC(=O)NCC1 | Not reached |
| 83 | Ethylene oxide | C1CO1 | Not reached |
| 84 | Diethanolamine | C(CO)NCCO | Not reached |
| 85 | Dimethylether | COC | Reached in both 1 post-PKS and 2 post-PKS steps |
| 86 | Propane | CCC | Reached in both 1 post-PKS and 2 post-PKS steps |
| 87 | Ethene | C=C | Reached in 2 post-PKS steps only |
| 88 | Propylene Oxide | CC1CO1 | Reached in 2 post-PKS steps only |

| | | | |
|---|---|---|---|
| 89 | Dihydrofuranone | C1C(COC1=O)O | Reached simply with PKSs |
| 90 | Ethyl acetate | CCOC(=O)C | Reached in 2 post-PKS steps only |
| 91 | Geraniol | CC(=CCCC(=CCO)C)C | Not reached |
| 92 | 2_Furanmethanol | C1=COC(=C1)CO | Not reached |
| 93 | Furfural | C1=COC(=C1)C=O | Not reached |
| 94 | Geranial | CC(=CCCC(=CC=O)C)C | Not reached |
| 95 | Benzamine | C1=CC=C(C=C1)N | Reached in 2 post-PKS steps only |
| 96 | Cyanuric acid | C1(=O)NC(=O)NC(=O)N1 | Not reached |
| 97 | Terephtalic_acid | COC(=O)C1=CC=C(C=C1)C(=O)O | Reached in 2 post-PKS steps only |
| 98 | Resorcinol | C1=CC(=CC(=C1)O)O | Reached in 2 post-PKS steps only |
| 99 | 3_Methylphenol | OC1=CC(C)=CC=C1 | Reached in 2 post-PKS steps only |
| 100 | Cresol | OC1=CC=C(C)C=C1 | Reached in 2 post-PKS steps only |
| 101 | Melamine | C1(=NC(=NC(=N1)N)N)N | Not reached |
| 102 | 9,10-anthraquinone | C1=CC=C2C(=C1)C(=O)C3=CC=CC=C3C2=O | Not reached |
| 103 | Styrene | C=CC1=CC=CC=C1 | Reached in both 1 post-PKS and 2 post-PKS steps |
| 104 | 14_Dimethyl_Benzene | CC1=CC=C(C=C1)C | Reached in 2 post-PKS steps only |
| 105 | Terpinolene | CC1=CCC(=C(C)C)CC1 | Not reached |
| 106 | 1_Naphthalenol | C1=CC=C2C(=C1)C=CC=C2O | Not reached |
| 107 | Coumarin | C1=CC=C2C(=C1)C=CC(=O)O2 | Not reached |
| 108 | 12_Benzenediol | C1=CC=C(C(=C1)O)O | Reached in 2 post-PKS steps only |
| 109 | Naphthalene | C1=CC=C2C=CC=CC2=C1 | Not reached |
| 110 | 13_Dimethyl_Benzene | CC1=CC(=CC=C1)C | Reached in 2 post-PKS steps only |
| 111 | Acetic_acid_phenylmethyl_ester | CC(=O)OCC1=CC=CC=C1 | Reached in 2 post-PKS steps only |
| 112 | Benzeneethanol | C1=CC=C(C=C1)CCO | Reached in both 1 post-PKS and 2 post-PKS steps |
| 113 | Bisphenol_ A | CC(C)(C1=CC=C(C=C1)O)C2=CC=C(C=C2)O | Not reached |
| 114 | 4_Ethylphenol | CCC1=CC=C(C=C1)O | Reached in 2 post-PKS steps only |
| 115 | Phenol | C1=CC=C(C=C1)O | Reached in both 1 post-PKS and 2 post-PKS steps |
| 116 | Caffeine | CN1C=NC2=C1C(=O)N(C(=O)N2C)C | Not reached |
| 117 | 2-methoxyphenol | COC1=CC=CC=C1O | Not reached |
| 118 | 3_Phenyl_2_propenal | C1=CC=C(C=C1)C=CC=O | Reached in both 1 post-PKS and 2 post-PKS steps |
| 119 | Benzenemethanol | C1=CC=C(C=C1)CO | Reached in both 1 post-PKS and 2 post-PKS steps |

| | | | |
|---|---|---|---|
| 120 | Toluene | CC1=CC=CC=C1 | Reached in both 1 post-PKS and 2 post-PKS steps |
| 121 | Benzene | C1=CC=CC=C1 | Reached in 2 post-PKS steps only |
| 122 | Salicylic_acid | C1=CC=C(C(=C1)C(=O)O)O | Reached in 2 post-PKS steps only |
| 123 | 4_Hydroxyacetophenone | CC(=O)C1=CC=C(C=C1)O | Reached in 2 post-PKS steps only |
| 124 | 24_Diaminotoluene | CC1=C(C=C(C=C1)N)N | Not reached |
| 125 | 2_Hydroxytoluene | CC1=CC=CC=C1O | Reached in 2 post-PKS steps only |
| 126 | 2-Methylnaphthalene | CC1=CC=C(C=CC=C2)C2=C1 | Not reached |
| 127 | Estragole | COC1=CC=C(C=C1)CC=C | Not reached |
| 128 | Biuret | C(=O)(N)NC(=O)N | Not reached |
| 129 | 1,2-Dimethylbenzene | CC1=CC=CC=C1C | Reached in 2 post-PKS steps only |
| 130 | 1,4-Benzenediol | C1=CC(=CC=C1O)O | Reached in 2 post-PKS steps only |
| 131 | Ethylbenzene | CCC1=CC=CC=C1 | Reached in both 1 post-PKS and 2 post-PKS steps |
| 132 | 2_Napthol | C1=CC=C2C=C(C=CC2=C1)O | Not reached |
| 133 | 3-Methylbenzoic acid | CC1=CC(=CC=C1)C(=O)O | Not reached |
| 134 | Phenylbenzene | C1=CC=C(C=C1)C2=CC=CC=C2 | Not reached |
| 135 | Benzaldehyde | C1=CC=C(C=C1)C=O | Reached in both 1 post-PKS and 2 post-PKS steps |
| 136 | alpha-Tocopherol | CC1=C(C2=C(CCC(O2)(C)CCCC(C)CCCC(C)CCCC(C)C)C(=C1O)C)C | Not reached |
| 137 | 4-Hydroxybenzoic acid | C1=CC(=CC=C1C(=O)O)O | Reached in 2 post-PKS steps only |
| 138 | 2-Aminophenol | C1=CC=C(C(=C1)N)O | Not reached |
| **139** | **2-Phenylphenol** | **C1=CC=C(C=C1)C2=CC=CC=C2O** | **Reached in 2 post-PKS steps only** |
| 140 | 2_Methylbenzoic acid | CC1=CC=CC=C1C(=O)O | Not reached |
| 141 | 4_Aminophenol | C1=CC(=CC=C1N)O | Reached in both 1 post-PKS and 2 post-PKS steps |
| 142 | Methyl_ester_benzoic_acid | COC(=O)C1=CC=CC=C1 | Reached in 2 post-PKS steps only |
| 143 | Styrene oxide | C1C(O1)C2=CC=CC=C2 | Not reached |
| 144 | Nicotinic acid | C1=CC(=CN=C1)C(=O)O | Not reached |
| 145 | Salicylaldehyde | C1=CC=C(C(=C1)C=O)O | Reached in 2 post-PKS steps only |
| 146 | Acetophenone | CC(=O)C1=CC=CC=C1 | Reached in both 1 post-PKS and 2 post-PKS steps |
| **147** | **O_methyl_anthranilate** | **COC(=O)C1=CC=CC=C1N** | **Not reached** |
| **148** | **Methyl salicylate** | **COC(=O)C1=CC=CC=C1O** | **Not reached** |
| **149** | **Benzyl benzoate** | **C1=CC=C(C=C1)COC(=O)C2=CC=CC=C2** | **Not reached** |
| **150** | **Nicotinamide** | **C1=CC(=CN=C1)C(=O)N** | **Not reached** |

| | | | |
|---|---|---|---|
| 151 | 1,3-Dimethylxanthine | CN1C2=C(C(=O)N(C1=O)C)NC=N2 | Not reached |
| 152 | p-Toluic acid | CC1=CC=C(C=C1)C(=O)O | Not reached |
| 153 | Phthalate | C1=CC=C(C(=C1)C(=O)O)C(=O)O | Reached in 2 post-PKS steps only |
| 154 | Benzoic acid | C1=CC=C(C=C1)C(=O)O | Reached in both 1 post-PKS and 2 post-PKS steps |
| 155 | Vanillin | COC1=C(C=CC(=C1)C=O)O | Not reached |

**SI Table 4. Biomanufacturing candidates from Wu et. al. not used in reporting BioPKS Pipeline's performance.**

| # | Name | SMILES | Synthesis |
|---|---|---|---|
| 1 | Urea | C1 metabolite | Not used for validation |
| 2 | Methanamine | C1 metabolite | Not used for validation |
| 3 | Hydrocyanic acid | C1 metabolite | Not used for validation |
| 4 | Dichloromethane | C1 metabolite | Not used for validation |
| 5 | Chloromethane | C1 metabolite | Not used for validation |
| 6 | Methanol | C1 metabolite | Not used for validation |
| 7 | Methanethiol | C1 metabolite | Not used for validation |
| 8 | Formic acid | C1 metabolite | Not used for validation |
| 9 | Formamide | C1 metabolite | Not used for validation |
| 10 | Carbon disulfide | C1 metabolite | Not used for validation |
| 11 | Formaldehyde | C1 metabolite | Not used for validation |
| 12 | Methanesulfonic acid | C1 metabolite | Not used for validation |
| 13 | Carbon monoxide | C1 metabolite | Not used for validation |
| 14 | Carbon dioxide | C1 metabolite | Not used for validation |
| 15 | Nitrobenzene | Charged species | Not used for validation |
| 16 | Nitroethane | Charged species | Not used for validation |
| 17 | 2-Nitrophenol | Charged species | Not used for validation |
| 18 | 4-Nitrophenol | Charged species | Not used for validation |
| 19 | 1-Methyl-2,4-dinitrobenzene | Charged species | Not used for validation |
| 20 | Trichloroethane | Chlorinated species | Not used for validation |
| 21 | 1,2-Dichloroethane | Chlorinated species | Not used for validation |
| 22 | Tetrachloroethane | Chlorinated species | Not used for validation |
| 23 | Chloroethane | Chlorinated species | Not used for validation |

| | | | |
|---|---|---|---|
| 24 | 4-Aminobenzenesulfonic acid | Sulfonated species | Not used for validation |
| 25 | Chlorobenzene | Chlorinated species | Not used for validation |
| 26 | Chloroacetic acid | Chlorinated species | Not used for validation |
| 27 | 4-Methylbenzenesulfonic acid | Sulfonated species | Not used for validation |
| 28 | Bisulfinylmethane | Sulfonated species | Not used for validation |
| 29 | Thiobismethane | Sulfonated species | Not used for validation |
| 30 | 1,3,5-Triazine-2,4-diamine, 6-chloro-N-ethyl-N'-(1-methylethyl) | Sulfonated species | Not used for validation |
| 31 | 1,4-Dichlorobenzene | Chlorinated species | Not used for validation |
| 32 | Trichloroacetyldehyde | Chlorinated species | Not used for validation |
| 33 | 2,4-Dichlorophenoxyacetic acid | Chlorinated species | Not used for validation |
| 34 | 4-Chlorobenzamine | Chlorinated species | Not used for validation |
| 35 | 2,5-Dichlorophenol | Chlorinated species | Not used for validation |
| 36 | 2,4-Dichlorophenol | Chlorinated species | Not used for validation |
| 37 | Dichloroacetyl chloride | Chlorinated species | Not used for validation |
| 38 | 1,1-Dichloroethane | Chlorinated species | Not used for validation |
| 39 | 1,2,4-Trichlorobenzene | Chlorinated species | Not used for validation |
| 40 | 1,1,1-Trichloroethane | Chlorinated species | Not used for validation |
| 41 | Epichlorohydrin | Chlorinated species | Not used for validation |

**SI Table 5. Runtime information and statistics for DORAnet.**

| Name | Reaction type | Starter | 1 generation runtime | 2 generation runtime |
|---|---|---|---|---|
| 1,5-Pentanediol | A + A; A + helper | OCCCCCO | 00:00:43 | 00:48:31 |
| Glucose | A + A; A + helper | OCC1OC(O)C(O)C(O)C1O | 00:01:59 | 01:45:20 |
| Phenol | A + A; A + helper | Oc1ccccc1 | 00:00:39 | 00:10:48 |
